# DAXX governs the silencing of LINE1 during spermatogenesis in mice

**DOI:** 10.1101/2025.07.24.666258

**Authors:** Zejia Li, Chaoyang Xiong, Jin Shen, Chen Tan, Dupeng Ma, Jun Chen, Rong Liu, Qi Sun, Weihao Gong, Wenming Yuan, Mengya Huang, Li Huang, Yueqiu Tan, Guohong Li, Mengcheng Luo

## Abstract

The suppression of transposable elements in germline is critical for safeguarding fertility. Here we identify that the DAXX, a potent transcription repressor and an H3.3 chaperone, is indispensable for transposable elements silencing during spermatogenesis. Male mice with conditional knockout of *Daxx* in germ cells mediated by *Ddx4*-cre display delayed meiotic progression, deformed and reduced production of sperms, and an age-dependent decline in fertility. The ablation of DAXX results in activation of evolutionarily young LINE1 and ERVs subfamilies. Mechanistically, DAXX interacts with DMAP1 to maintain DNA methylation of LINE1 in meiotic cells. In summary, this study identifies DAXX as a previously unknown regulator of spermatogenesis and demonstrates that it functions as an epigenetic regulator to maintain the silencing the young LINE1 by DNA methylation.

## INTRODUCTION

DAXX (death associated protein 6), which was initially identified as an adaptor protein associated with the intracellular death domain of the apoptosis receptor Fas, is a highly conserved protein and plays diverse roles in fundamental biological pathways such as Fas-triggered apoptosis, tumor development, antiviral defense, transcription, cell cycle regulation, proliferation and multiple physiological functions ^1–3^. Biochemical and cell-based studies have demonstrated that DAXX serves as a specific chaperone for histone H3.3 that collaborates with the ATRX (alpha-thalassemia/mental retardation X-linked) to drive nucleosome assembly through replication-independent mechanisms. Furthermore, DAXX and ATRX facilitate the deposition of H3.3 into repetitive genomic regions, including pericentromeric DNA, telomeres, and transposable elements ^3,4^.

While retrotransposons function as evolutionary catalysts facilitating genomic diversification, they concurrently pose risks to host genome integrity. This is particularly true in germ cells when epigenetic control is relaxed, allowing for extensive genome-wide reprogramming. In mammals, the activation of retrotransposons induces sterility typically ^5^. To combat these risks, mammalian germline deploys multilayered regulatory strategies to restrict retrotransposon activity during critical developmental windows, including transcriptional repression via DNA methylation and H3K9 methylation, post-transcriptional silencing mediated by PIWI-interacting RNAs (piRNAs) ^6–8^. LINE1, the most abundant type of retrotransposons in the mammalian genome, poses a significant risk for infertility, especially in mouse mutants that exhibit overactivated retrotransposons (LINE1 and/or IAP) in male germ cells ^5^. Studies have also demonstrated that RLTR10B sequences, such as those found in MMERVK10C—one of the youngest retrotransposons in the mouse genome, are robustly silenced during spermatogenesis by TRIM33, a ubiquitin ligase that has E3 ubiquitin ligase activity for A-MYB and regulates its abundance for the transcription of any retrotransposon ^9^.

DNA methylation constitutes the principal epigenetic modification responsible for transcriptional repression of transposable elements (TEs) ^10^. DNMT3A (DNA Methyltransferase 3 Alpha) and DNMT3B (DNA Methyltransferase 3 Beta) mediate de novo DNA methylation, while DNMT1 (DNA Methyltransferase 1) maintains methylation profiles during DNA replication ^11–14^. Although the mechanisms of DNA de novo methylation have been under scrutiny ^15–18^, little is known about how the germ cells maintain DNA methylation to silence TEs persistently. The epigenetic reprogramming in primordial germ cells (PGCs) witnesses global methylation erasure, triggering transposon activation ^10^. This developmental process engages the piRNA biogenesis, where transposon-derived transcripts feed piRNA production ^19^. These piRNAs subsequently guide de novo methylation pattern at TEs promoters, initiating transcriptional repression via a molecular mechanism that remains incompletely resolved in mammals ^20–23^. However, how TEs activity is persistently suppressed and its methylation is maintained during the long process of spermatogenesis remains poorly understood. As ongoing research continues to uncover novel factors involved in the epigenetic silencing of retrotransposons in the germ cells, understanding these mechanisms may provide vital insights into fertility preservation and maintaining genome stability.

Here, we generate a germ-cell-specific *Daxx* knockout mouse model by crossing *Daxx*^flox/flox^ mice with *Ddx4*-Cre transgenic mice as global DAXX deletion is embryonically lethal ^24^, and demonstrates that DAXX loss in germ cells results in delayed spermatogenesis, abnormal sperm morphology, and an age-dependent decline in male fertility. Comprehensive transcriptome profiling reveals that DAXX loss induces intensive gene dysregulation, resulting in upregulation of apoptosis and transcription related genes. Mechanically, DAXX cooperates with DMAP1 to enforce LINE1 DNA methylation and epigenetically silence LINE1 in meiotic cells.

## RESULTS

### DAXX is a nuclear protein and highly expressed in spermatocytes and early-stage spermatids

*Daxx* mRNA is ubiquitously expressed in various kinds of organs or tissues and particularly the highest in the testes of both humans and mice (Figure. S1A and S1C) ^25,26^, and single cell sequencing reported relatively higher levels in germ cells undergoing meiotic prophase I and early spermatids (Figure. S1B and S1D) ^27,28^. Although *Daxx* mRNA was detected across various organs or tissues (Figure. S1), DAXX protein was predominantly present in the testes of adult mice (Figure. 1A). Immunoblotting of total protein extracts from wildtype (WT) mouse testes at different ages [from postnatal day 3 (P3D) to postnatal day 35 (P35D)] revealed that DAXX protein begins to express at very low level on P3D, increases notably since P14D, and peaks on P21D (Figure. 1B). To further explore the biological function of DAXX in spermatogenesis, we isolated various developmental stages of male germ cells using fluorescence-activated cell sorting (FACS) ^29,30^. Western blot analysis showed that DAXX is highly expressed from the primary spermatocytes to the round spermatids (RS) stage, but has extremely low level in elongating spermatids and mature sperms (Figure. 1C). To map the its intracellular compartmentalization, we performed nuclear and cytoplasmic fractionation of mouse testis extracts, and western blot analysis revealed that DAXX was present exclusively in the nucleoplasmic fraction, and it exhibited limited binding capacity to chromatin, but it was almost absent or undetectable in cytoplasm (Figure. 1D). These findings suggest that DAXX could be associated with nuclear events in spermatocytes and early-stage spermatids.

**Figure 1.**
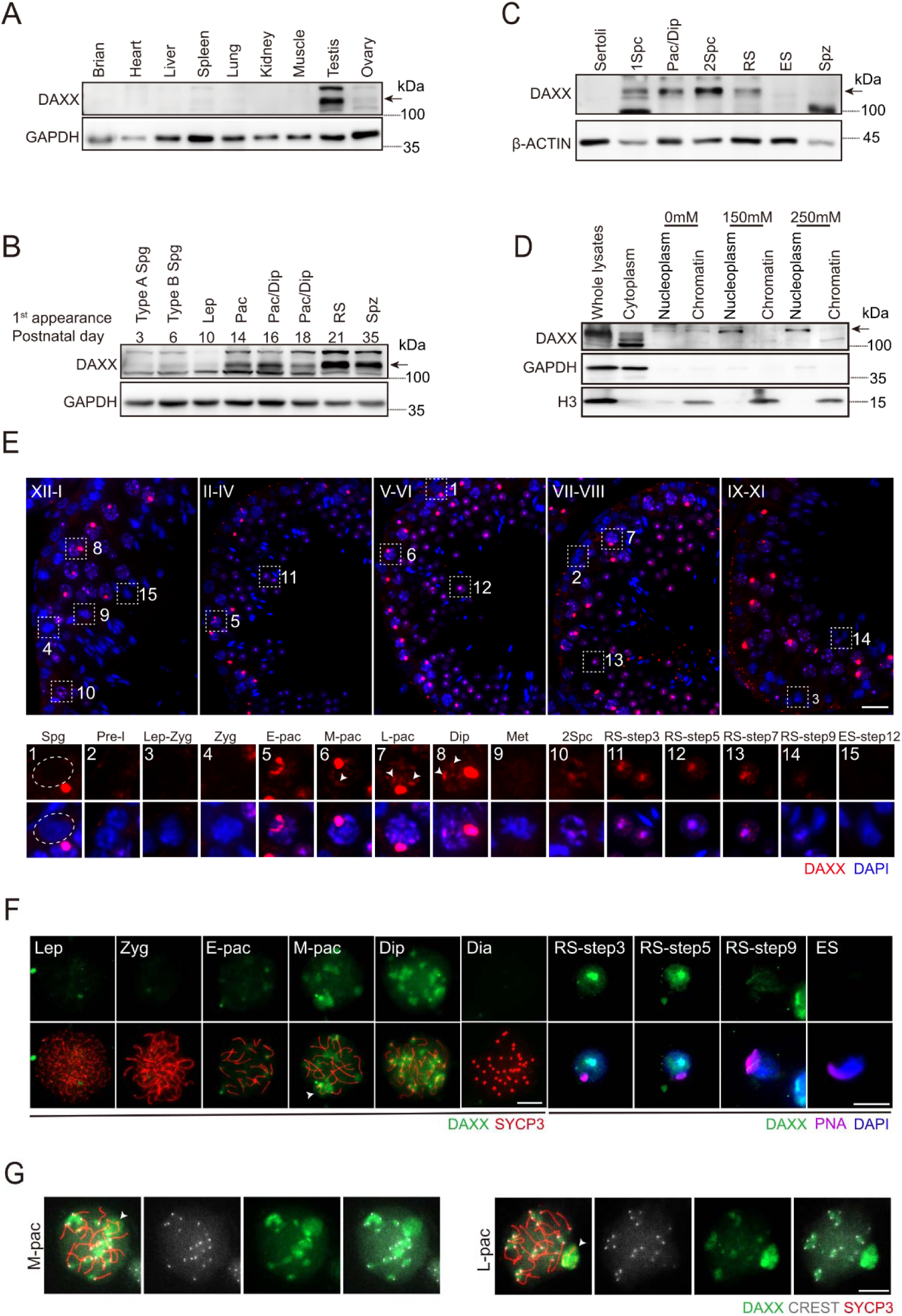
The dynamic expression and localization of DAXX. (A) DAXX expression profile in adult mouse tissues/organs detected by western blot. GAPDH serves as the loading control. The arrow indicates the specific DAXX band. (B) Temporal expression profile of DAXX in mouse testes detected by western blot. GAPDH serves as the loading control. The arrow indicates the specific DAXX band. (C) DAXX expression in isolated male germ cells and Sertoli cells detected by western blot, β-ACTIN serves as the loading control. The arrow indicates the specific DAXX band. (D) DAXX expression in the cytoplasmic, nucleoplasm and chromatin of adult WT testes detected by western blot. GAPDH serves as a marker for cytoplasmic component, and histone H3 serves as a marker for chromatin component. The arrow indicates the specific DAXX band. (E) Representative images of immunofluorescence staining of DAXX (red) on testicular sections from WT adult mice. Scale bar, 20 μm. (F) Chromosome spreads of spermatocytes from adult WT testes were immunostained for DAXX (green), SYCP3 (red), PNA (purple) and DAPI (blue). The white arrowhead indicates the XY body. Scale bar, 10 μm. (G) Immunofluorescence staining of DAXX (green), SYCP3 (red) and CREST (grey) on chromosome spreads of spermatocytes from adult WT testes. The white arrowhead indicates the XY body. Scale bar, 10 μm. Abbreviation in B, C, E and F: Spg: spermatogonia; 1Spc: primary spermatocyte; 2Spc: secondary spermatocyte; Pre-l: preleptotene; L: leptotene; Z: zygotene; P: pachytene; E-pac: early pachytene; M-pac: middle pachytene; L-pac: late pachytene; Dip: diplotene; Dia: diakinesis; Met: meiotic metaphase I; RS: Round spermatid; ES, elongating spermatid; Spz: spermatozoa.

To identify the stage-specific role of DAXX, we examined its localization and expression in testicular sections from adult WT mice. Immunofluorescence (IF) analysis revealed that DAXX first appears at stages II– IV of the seminiferous epithelial cycle, showing prominent accumulation at the XY body in pachytene and diplotene spermatocytes. Notably, several intensely-stained DAXX blobs are observed in the nucleus from mid-pachytene to diplotene spermatocytes (Figure 1E, arrowheads). While DAXX signal persists strongly in diplotene spermatocytes, it disappears as cells progress into meiotic metaphase I and reappear in secondary spermatocytes and RS. No DAXX signal is detected in spermatogonia, leptotene/zygotene spermatocytes, and elongating/condensed spermatids (Figure 1E).

We next determined DAXX’s subcellular localization patterns via chromosome spread analysis. Precise substaging of meiosis was achieved by labeling the synaptonemal complex protein SYCP3, a critical component of the axial element. Consistent with the above results, we found that DAXX was expressed in the nuclear from spermatocytes to round spermatids, with pronounced enrichment at chromosome ends and the XY body of the pachytene and diplotene spermatocytes (Figure. 1F). To further refine the DAXX localization within the nucleus, we employed CREST staining to label the centromeres, and found marked aggregation of DAXX at the centromeres (Figure. 1G), suggesting a potential function association with these structures during meiosis. Together, these findings provide strong evidence that DAXX is not only present but also maybe involve in critical stages of germ cell development, particularly in meiotic chromosome dynamics. This distinct localization pattern at centromeres and the XY body highlights DAXX’s potential as a key regulatory player in spermatogenesis, underscoring its importance for maintaining male fertility.

### DAXX deletion leads to impaired spermatogenesis and age-dependent fertility loss in male mice

To investigate the role of DAXX in mammalian spermatogenesis, we developed a conditional knockout mouse model with specific deletion of *Daxx* in germ cells mediated by *Ddx4*-cre due to its global loss leads to embryonic lethality ^24^. In this model, exon 2 to 7 of *Daxx* is flanked by two loxP sites, which is then deleted by *Ddx4*-Cre recombinase in testis (*Daxx*^flox/−^, *Ddx4*-Cre mice, referred to as *Daxx*^cKO^) (Figure. 2A). The *Ddx4*-Cre transgene exhibits germ cells-restricted activation since embryonic day 15 (E15) and continues postnatally ^31^. PCR-based genotyping analyses of mouse tail confirmed the genotypes of *Daxx*^cKO^, *Daxx*^flox/+^ and *Daxx*^+/+^ mice (Figure 2B). Western blot assay indicated that DAXX protein level was significantly decreased in *Daxx*^cKO^ testes compared to the control testes (Figure. 2C). Immunofluorescence (IF) analysis also demonstrated depletion of DAXX protein within *Ddx4* positive germ cells in *Daxx*^cKO^ testes, while standard expression patterns were maintained in control littermates (Figure 2D).These collectively indicate that DAXX was successfully knocked out in male germ cells.

**Figure 2.**
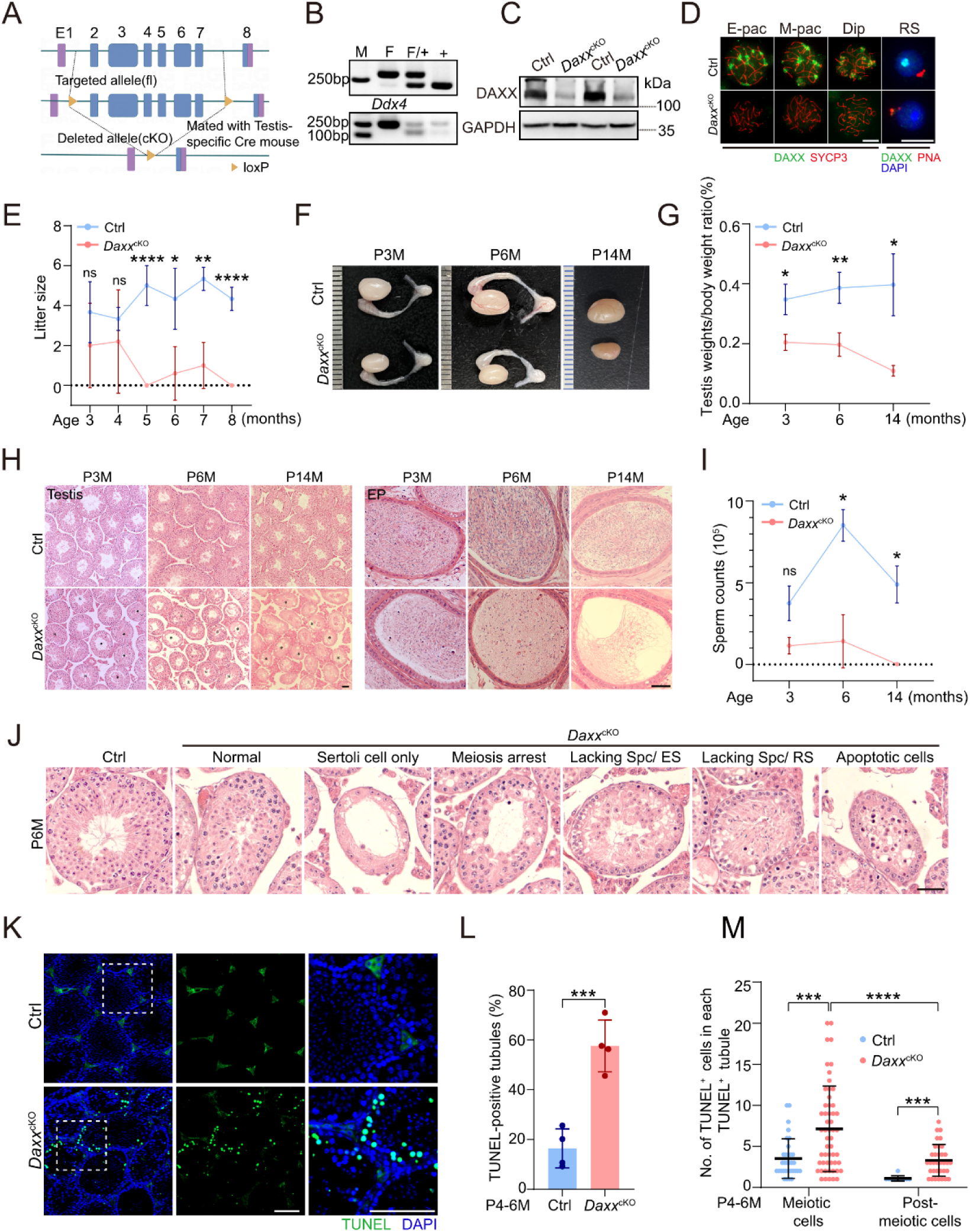
DAXX depletion leads to impaired spermatogenesis and age-dependent fertility loss in male mice. (A) Schematic representation of the CRISPR-Cas9 genome editing system to generate the *Daxx*-flox mice and mating strategy. (B) Agarose gel electrophoresis of representative genotyping PCR results of F (flox), *+* (WT) and *Ddx4-cre* using the primer sets in Table S1. The PCR product of WT allele (“+”) is 195 bp, the product of floxed allele (“F”) is 280 bp and the product of *Ddx4-cre* allele is 240 bp. (C) DAXX was deleted in adult *Daxx*^cKO^ testes compared to control detected by western blot. GAPDH serves as the loading control. (D) DAXX was deleted in adult *Daxx*^cKO^ testes compared to control detected by Immunofluorescence staining of chromosome spreads of spermatocytes. E-pac: early pachytene; M-pac: middle pachytene; L-pac: late pachytene; Dip: diplotene; RS: Round spermatid. Scale bar, 10 μm. (E) Litter size per female mated with WT or *Daxx*^cKO^ male mice since 2 months of age during 6-month breeding trial. (F) Testis morphology of WT and *Daxx*^cKO^ mice at 3, 6 and 14 months of age. (G) Ratio of testis/body weight of control and *Daxx*^cKO^ mice at 3, 6 and 14 months of age. (H) Hematoxylin and eosin (H&E) staining of testis and cauda epididymis sections from control and *Daxx*^cKO^ mice. “N” indicates normal tubules, “IZ” indicates disordered and sparse tubules, “#” indicates meiotic defect, “*” indicates Sertoli cells only, respectively. Scale bar, 50 μm. (I) Quantification of sperm counts in control and *Daxx*^cKO^ male mice. (J) Histology analysis of testes from 6-month-old control and *Daxx*^cKO^ mice. Scale bar, 50 μm. (K) TUNEL staining of testicular sections from control and *Daxx*^cKO^ mice. Scale bar, 100 μm. (L) The plot shows the percentage of testicular cross-sections containing apoptotic cells in control and *Daxx*^cKO^ testes. ***p < 0.001 using two-tailed Student’s unpaired t-test. (M) The plot shows the number of TUNEL-positive cells per TUNEL-positive tubular cross-section in control and *Daxx*^cKO^ testes. ns, not significant, *p < 0.05, **p < 0.01, ****p < 0.0001 using Multiple Mann-Whitney tests.

Subsequently, fertility tests revealed that it was almost normal for *Daxx*^cKO^ males up to 4 months old as they produced comparable numbers of offspring when compared with the controls (Figure. 2E). However, WT females mated with *Daxx*^cKO^ males at mid-adulthood (5 to 6 months of age) and even later had much smaller litter size (Figure. 2E). The testes from 3, 6 and 14-month-old *Daxx*^cKO^ mice were significantly smaller than those of the controls (Figure. 2F and 2G). We speculate that DAXX loss may cause an age-dependent decline in male fertility, so we performed histological analysis of their testes and epididymis in different age groups. In contrast to the seminiferous epithelium of control mice, which had the typical arrangement of spermatogenic cells including spermatogonia, spermatocytes, round spermatids, and elongating spermatids from the basal membrane to the lumen, *Daxx*^cKO^ mice exhibited an age-dependent loss of germ cells and extensive defects across the 12 stages of the seminiferous epithelium (Figure. 2H). We also quantified the sperm in the cauda epididymis and found that *Daxx*^cKO^ mice had significantly fewer sperm than control mice (Figure. 2I). The testes from 6-month-old *Daxx*^cKO^ male mice displayed progressive germ cell depletion along spermatogenic stages, initiating from early germ cells (spermatogonia) to late germ cells (elongated spermatids) (Figure. 2J).An age-dependent increase of the percent of abnormal tubules was observed, while the population of spermatogonia progressively decreases in *Daxx*^cKO^ male mice (Figure. S2). Furthermore, we performed a TUNEL assay and found that the number of TUNEL-positive tubules and cells in the seminiferous tubules of *Daxx*^cKO^ mice was significantly higher than that in control males (Figure. 2K-2M), including more apoptotic spermatocytes and post-meiotic cells.

### DAXX loss does not affect meiotic chromosomal behaviors

Given that extensive apoptotic spermatocytes and post-meiotic cells were observed in *Daxx*^cKO^ testis, we first hypothesized that DAXX loss could result in meiotic defects. To explore events leading to apoptotic meiocytes in *Daxx*^cKO^ male mice, we examined DSB formation and synapsis of homologous chromosomes by IF staining for SYCP3 and SYCE1 (which are essential structural components of the synaptonemal complex (SC)) or γH2AX (which participates in DSB repair and is robustly expressed in the nucleus from leptotene to zygotene stages, and also enriched in the sex body (XY body) during the pachytene and diplotene stages). However, we found no difference in SYCP3 and SYCE1 or γH2AX signal between Ctrl and *Daxx*^cKO^ spermatocytes (Figure. S3A and S3B), suggesting that synapsis of homologous chromosomes and DSB formation and repair both progressed normally during these stages in *Daxx*^cKO^ spermatocytes. Furthermore, primary spermatocytes from *Daxx*^cKO^ mice progressed normally through leptotene, zygotene, pachytene, and diplotene stages with meiotic substage distributions comparable to controls, indicating assembly and disassembly of the synaptonemal complex proceeded normally in *Daxx*^cKO^ spermatocytes (Figure. S3C). During meiotic DSB repair and recombination, RPA2 and MLH1/3 mediate strand binding and crossover formation, respectively. Immunostaining analysis revealed comparable numbers of RPA2 between *Daxx*^cKO^ and control spermatocytes (Figure. S3D and S3E). MLH1 foci quantification also showed no significant difference between *Daxx*^cKO^ spermatocytes versus controls (Figure. S3F and S3G), suggesting that crossover recombination was intact in *Daxx*^cKO^ spermatocytes. CENPA (Centromere Protein A) is the core histone variant of centromeric chromatin, which governs faithful chromosome segregation. *Daxx*^cKO^ spermatocytes exhibited CENPA localization patterns indistinguishable from controls (Figure. S3H). These results indicate that DAXX loss does not affect chromosomal behavior in primary spermatocytes during meiosis.

### Loss of DAXX causes longer Manchette MTs and damaged acrosomes

To further investigate the spermiogenic defects in *Daxx*^cKO^ mice based on our observation of increased apoptotic spermatids in the seminiferous epithelium and reduced sperm production, we performed double immunofluorescence staining of peanut agglutinin (PNA, a sperm acrosome marker), γH2AX (for staging of seminiferous tubules) and DAPI for labeling DNA on testicular sections (Figure.3A). Notably, the transition of round spermatids into elongating spermatids was significantly impaired at stage X-XI in *Daxx*^cKO^ spermatids. Nuclear dysmorphology (stage X-XI, step 10-11) with loss of cephalic hook formation and dorsal angulation acrosomal extension defects concomitant with dorsoventral fin structure depletion, and the subsequent elongating spermatids at step 12 exhibited morphological abnormalities. It is noteworthy that compared to the controls, haploid spermatids at steps 1–16 were obviously reduced in the seminiferous tubules of *Daxx*^cKO^ testes (Figure. 3B). Using FACS, we also compared the proportions of specific germ cell populations between 4-month-old control and *Daxx*^cKO^ mice (Figure. S4). Consistent with the results of PNA staining, we observed a significant reduction in the haploid cells in the *Daxx*^cKO^ males (46.3% in controls *v.s.* 37.6% in *Daxx*^cKO^) (Figure. S4A and S4B).

**Figure 3.**
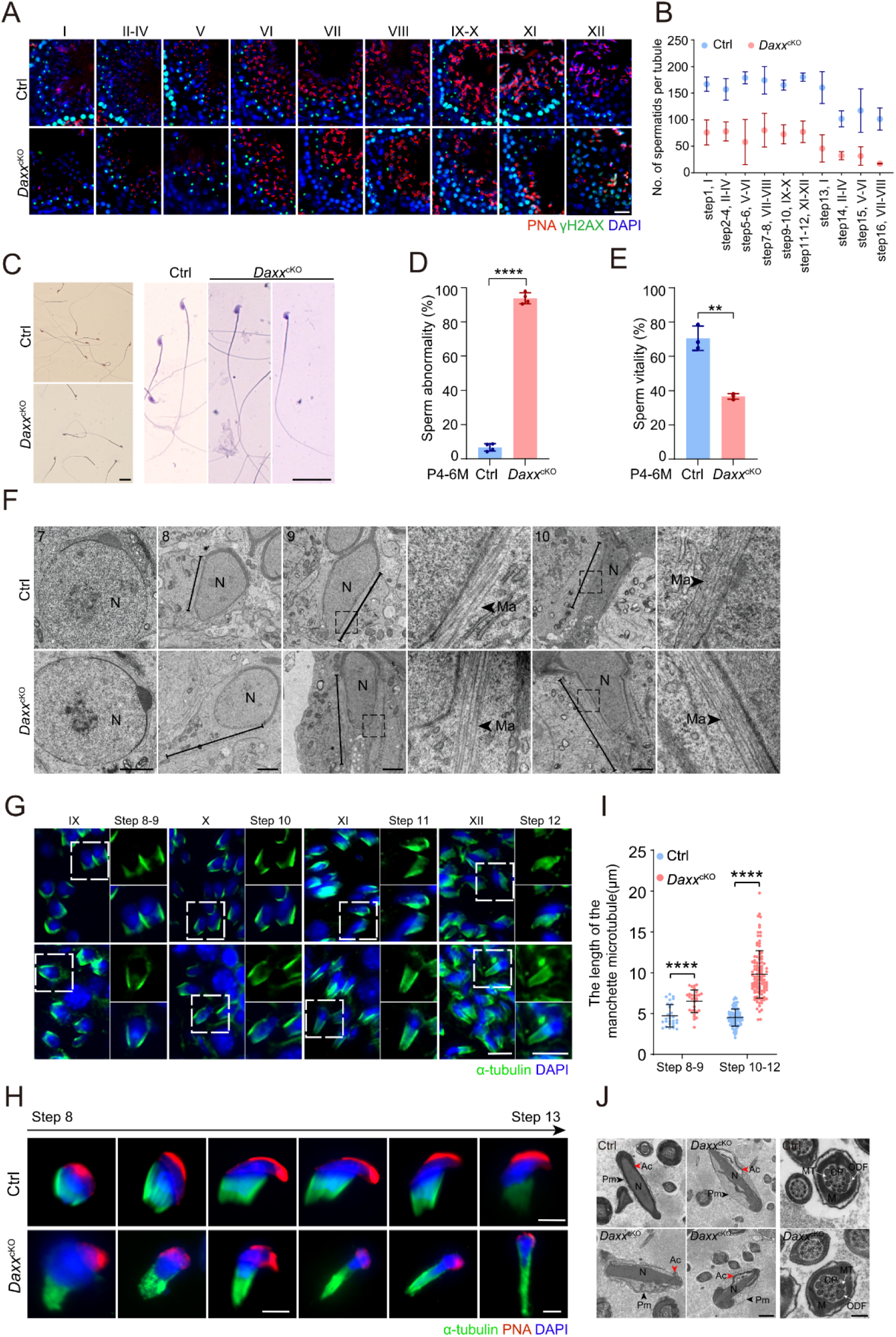
DAXX loss causes abnormally longer manchette MTs and acrosomal defects. (A) Representative images of immunofluorescence staining of γ-H2AX (green), PNA (red), and DAPI (blue) on stage-specific seminiferous tubule cross-sections from control and *Daxx*^cKO^ mice. The co-stained γH2AX was used for staging of spermatocytes. Scale bar, 20 μm. (B) Quantitative analysis revealed significant reduced spermatids in *Daxx*^cKO^ testes. N=3. (C) Diff Quick-stained epididymal sperm from adult control and *Daxx*^cKO^ littermates. *Daxx*^cKO^ sperm exhibit abnormal nuclear morphology. Scale bars, 20 μm. (D) Quantitative of epididymal sperm with abnormal morphology between control and *Daxx*^cKO^ littermates. \*\*\*\**p* < 0.0001 using two-tailed Student’s unpaired *t*-test. (E) Quantitative of epididymal sperm motility between adult control and *Daxx*^cKO^ mice. \*\**p* < 0.01 using two-tailed Student’s unpaired *t*-test. (F) The ultrastructure of step 8 to 10 spermatids from control and *Daxx*^cKO^ mouse testes. N, nucleus. Ma, manchette. Scale bars, 2 μm. (G) Immunofluorescence staining of α-tubulin (red) for manchette MTs and DAPI (blue) for nucleus in testicular sections at different stages from control and *Daxx*^cKO^ mice. Scale bars, 10 μm. (H) Comparison of manchette MTs length between control and *Daxx*^cKO^ in isolated elongating spermatids are shown in progressive steps during head elongation. α-tubulin staining (green) for manchette MTs, PNA staining (red) for the acrosome and DAPI (blue) staining for the nucleus. Scale bars, 5 μm. (I) Quantification of manchette MTs length in isolated spermatids from control and *Daxx*^cKO^ mice. Control: n = 27 for steps 8 to 9; n = 85 for step 10-12; *Daxx*^cKO^: n = 30 for steps 8 to 9; n = 108 for step 10-12. \*\*\*\**p* < 0.0001 using Multiple Mann-Whitney tests. (J) TEM revealed acrosomal detachment and plasma membrane discontinuity in *Daxx*^cKO^ spermatozoa. The red arrowhead indicates the acrosome. The black arrowhead indicates the plasma membrane. N, nucleus; Ac, acrosome; Pm, plasma membrane. Scale bars, 1 μm. MT, microtubules; M, mitochondrial sheath; CP, central microtubules; ODF, outer dense fibers. Scale bars, 200 nm.

We then assessed the sperm parameters and found that the proportion of mature sperm with head malformations in the epididymis of *Daxx*^cKO^ mice was higher than that in control mice (Figure. 3C and 3D). Additionally, computer-assisted sperm analysis (CASA) revealed a reduction in sperm motility in *Daxx*^cKO^ mice (Figure. 3E). Sperm head malformations frequently correlate with damaged acrosomal formation and manchette structure ^32^. These structures play pivotal roles in the elongation of spermatids through their functional synergy ^33,34^. The acrosome is a special kind of organelle with a cap-like structure that covers the head of the spermatid ^35,36^. The manchette is a typical non-centrosomal microtubule-enriched structure ^37^. To analyze the manchette microtubules (MTs) comprehensively, we employed transmission electron microscopy (TEM) imaging of testis sections, along with immunostaining (Figure. 3F-3H). The results showed that while *Daxx*^cKO^ spermatids maintained structurally normal manchette MTs up to step 7, the manchette MTs surrounding the nucleus became abnormally elongated during steps 8 to 10 of spermatid development (Figure. 3F). Additionally, abnormally longer manchette MTs were also showed in frozen testis sections of *Daxx*^cKO^ mice (Figure. 3G). To eliminate the potential observation bias from the different direction of the section, we performed immunofluorescence staining of single spermatids (Figure. 3H). Consistent with the above findings, the MTs of the manchette in elongating spermatids from *Daxx*^cKO^ mice continued to lengthen, resulting in a longer and thinner nucleus that lacked the characteristic hook shape as spermiogenesis progressed when compared with those from the controls (Figure. 3H and 3I). PNA signals also exhibited a nonuniform distribution in elongating spermatids in *Daxx*^cKO^ mice (Figure. 3H). To elucidate the acrosomal damaged, ultrastructural profiling of spermatozoa was performed through TEM. In contrast to spermatozoa from control mice, which exhibited long oval shapes with their acrosomes attached to the nuclei, the acrosome of *Daxx*^cKO^ males showed dissociation from the nuclear envelope, presenting as a shedding and pleating appearance but normal ‘9+2’ structure (Figure. 3J). These findings suggest that DAXX is essential for the acrosome formation and the organization of the manchette MTs network during spermiogenesis.

Given DAXX’s role as a molecular chaperone for histone H3.3, it is likely that it participates in the histone-to-protamine replacement during sperm maturation. To investigate this, we assessed nuclear condensation by staining for transition protein 1 (TNP1) and protamine 1 (PRM1) and observed that a marked reduction in the fluorescence intensity of TNP1 and PRM1 in the testicular sections of *Daxx*^cKO^ mice when compared with the controls (Figure. S5A and S5B). Additionally, western blot analysis revealed a significant decrease of PRM1 protein expression in the *Daxx*^cKO^ testes (Figure. S5C). To eliminate the possibility that the reduced PRM1 levels were resulted from a reduction in sperm count (as indicated in Figure. 2I), we collected mature sperm from the cauda epididymis and performed western blot for evaluating PRM1’s abundance. The results still reported a significant reduction in PRM1 levels of mature sperm in *Daxx*^cKO^ mice (Figure. S5D). We also quantified the levels of histones H3 in the testicular section and found that histone H3 has been replaced by PRM1 in tubules from stages XII to I (Figure. S5E). These findings suggest although there was a reduction in PRM1 abundance due to sperm defects, the process of histone-to-protamine replacement remained unaffected.

### The first wave of spermatogenesis was delayed in *Daxx*^cKO^ mice

In the first wave of spermatogenesis, synchronized germ cell progress through the seminiferous epithelium with precise patterning - exhibiting stage-specific emergence from primordial germ cells to terminally differentiated spermatozoa, and this is different from the adult ^38^. To better understand the role of DAXX in the first wave of spermatogenesis, we measured the testis weight of mice at various time points after birth through adulthood. Testis weight increased progressively with age before adulthood across both genotypes. However, a significant reduction in testis weight was observed in *Daxx*^cKO^ mice starting from P25D (Figure. 4A). To identify the exact phases of spermatogenesis caused by DAXX deletion, we conducted the H&E analysis of the testes of juvenile mice from P10D through P40D. We found that there were comparable numbers of spermatocytes in WT and *Daxx*^cKO^ testes at P10D (Figure. 4B), indicating that DAXX loss didn’t affect meiotic initiation. At P14D, the seminiferous tubules contained many early pachytene spermatocytes in the control testes, while the number of tubules containing pachytene spermatocytes was much lower in *Daxx*^cKO^ testes (Figure. 4B). By P18D, control tubules showed accumulation of late pachytene/diplotene spermatocytes, whereas *Daxx*^cKO^ displayed a significant reduction of these populations (Figure. 4B). At P25D, control testes exhibited round spermatid production, while *Daxx*^cKO^ showed a dramatic absence of post-meiotic cells (Figure. 4B). In contrast to controls where spermiogenesis culminated in elongated spermatids, *Daxx*^cKO^ mice at P40D exhibited the reduction or even absence of elongated spermatids, demonstrating slowed progression through meiosis (Figure. 4B).

**Figure 4.**
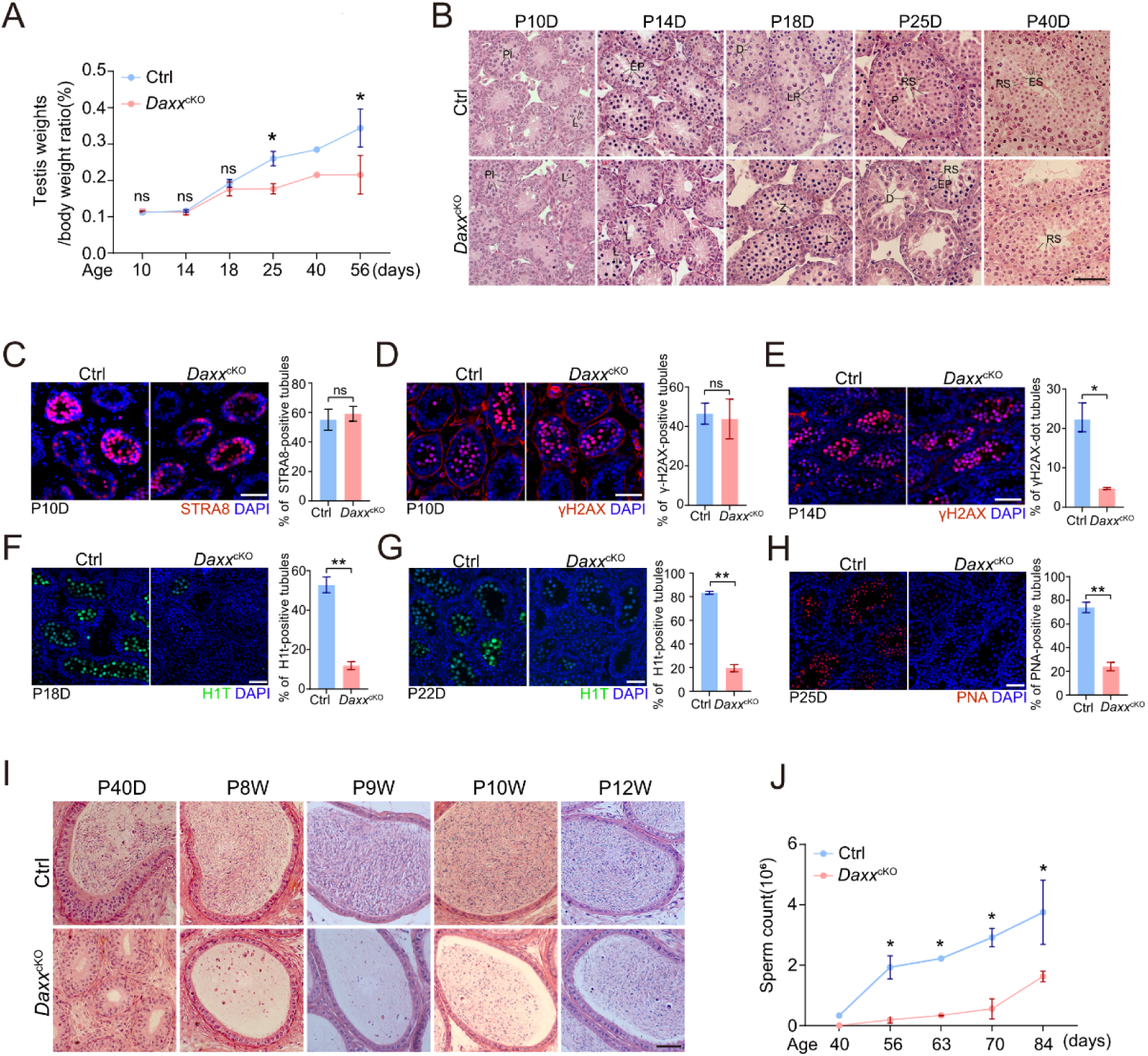
The first wave of spermatogenesis is delayed in *Daxx*^cKO^ mice. (A) Ratio of testis weight/body weight of control and *Daxx*^cKO^ males during the first wave of spermatogenesis. (B) Representative images of H&E-stained testicular cross-sections of control and *Daxx*^cKO^ mice from P10D through P40D. Pl: pre-leptotene; L: leptotene; Z: zygotene; EP: early-pachytene; LP: late-pachytene; D: diplotene; RS: round spermatids; ES: elongating spermatids. Scale bar, 50 μm. (C) Immunofluorescence staining of STRA8 (red) and quantification of the STRA8-positive tubules in testicular cross-section from control and *Daxx*^cKO^ mice at P10D, N=2. Scale bar, 50 μm. (D) Immunofluorescence staining of γH2AX (red) and quantification of the γH2AX-positive tubules in testicular cross-section from control and *Daxx*^cKO^ mice at P10D, N=2. Scale bar, 50 μm. (E) Immunofluorescence staining of γH2AX (red) and quantification of the γH2AX-dot tubules in testicular cross-section from control and *Daxx*^cKO^ mice at P14D, N=2. Scale bar, 50 μm. (F) Immunofluorescence staining of H1T (green) and quantification of the H1T-positive tubules in testicular cross-section from control and *Daxx*^cKO^ mice at P18D, N=2. Scale bar, 50 μm. (G) Immunofluorescence staining of H1T (green) and quantification of the H1T-positive tubules in testicular cross-section from control and *Daxx*^cKO^ mice at P22D, N=2. Scale bar, 50 μm. (H) Immunofluorescence staining of PNA (red) and quantification of the PNA-positive tubules in testicular cross-section from control and *Daxx*^cKO^ mice at P25D, N=2. Scale bar, 50 μm. (I) Representative images of H&E-stained cauda epididymis from control and *Daxx*^cKO^ mice at P40D, postnatal weeks 8 (P8W), P9W, P10W, and P12W. Scale bar, 50 μm. (J) Sperm counts in control and *Daxx*^cKO^ male mice at different ages (from P40D to P84D). Data in C-H, Statistics: two-tailed Student’s unpaired *t*-test, ns, not significant, \**p* < 0.05, \*\**p* < 0.01.

Immunofluorescence staining of representative stages of the first wave of spermatogenesis by labeling various markers, including STRA8 (a retinoic acid induced gene that plays a vital role in meiotic entry), γH2AX (a marker for DNA DSBs and thus spermatocytes), H1T (a marker expressed in mid to late pachytene cells but not in early pachytene cells) ^39^, and PNA (a marker for sperm acrosome), further revealed the slower progression of the first wave of spermatogenesis in *Daxx*^cKO^ mice (Figure. 4C-H). Additionally, we also performed histological analysis of the cauda epididymis at postnatal weeks 6, 8, 9, 10 and 12 and found that sperm began to appear in the cauda epididymis of control mice at P40D, whereas sperms were not detected until postnatal weeks 8 in *Daxx*^cKO^ mice (Figure 4I and 4J). Collectively, these data provide clear evidences that the first wave of spermatogenesis is delayed in *Daxx*^cKO^ mice.

### DAXX loss alters the transcriptome of germ cells

The absence of DAXX may disrupt the expression of specific genes critical for spermatogenesis, leading to severe apoptosis of germ cells and abnormal sperm morphology. Then we performed genome-wide transcriptome analyses of isolated spermatocytes and RS and identified 60 significantly upregulated genes and 22 significantly downregulated genes in *Daxx*-deficient meiotic cells (Figure. 5A). In contrast, DAXX-deficient RS exhibited much greater changes in gene expression, that 2,144 genes were upregulated and 176 genes downregulated (Figure. 5B). The downregulated genes in both meiotic and post-meiotic cells were enriched in cell membrane components (Figure. 5D and 5F), while many upregulated genes were associated with RNA polymerase II-mediated transcription and apoptotic pathways (Figure. 5E and 5G). Notably, genes associated with actin cytoskeleton organization were downregulated in DAXX-deficient RS (Figure. 5F), which may be one of the reasons for the aforementioned Manchette MTs abnormalities observed in *Daxx*^cKO^ mice. A comparative analysis of meiotic and post-meiotic transcriptomes in control cells confirmed previous findings ^40,41^ that the transition from meiosis to post-meiosis involves extensive transcriptional reprogramming, in which thousands of genes either upregulated or downregulated, were organized into distinct clusters (Figure. S6A-S6C). DAXX deletion resulted in notable shifts in gene regulation during this transition. Specifically, quite genes were downregulated, while more genes were upregulated in *Daxx*^cKO^ cells, particularly those involved in transcription. However, the overall clustering patterns remained unaffected (Figure. S6C). Although gene dysregulation followed similar trends in both control and *Daxx*^cKO^ cells, the number of upregulated genes was larger in *Daxx*^cKO^ cells (Figure. S6A). Taken together, these findings highlight DAXX’s essential role in regulating the transcriptional landscape of germ cells and it has different effects on specific genes of different developmental processes during spermatogenesis.

**Figure 5.**
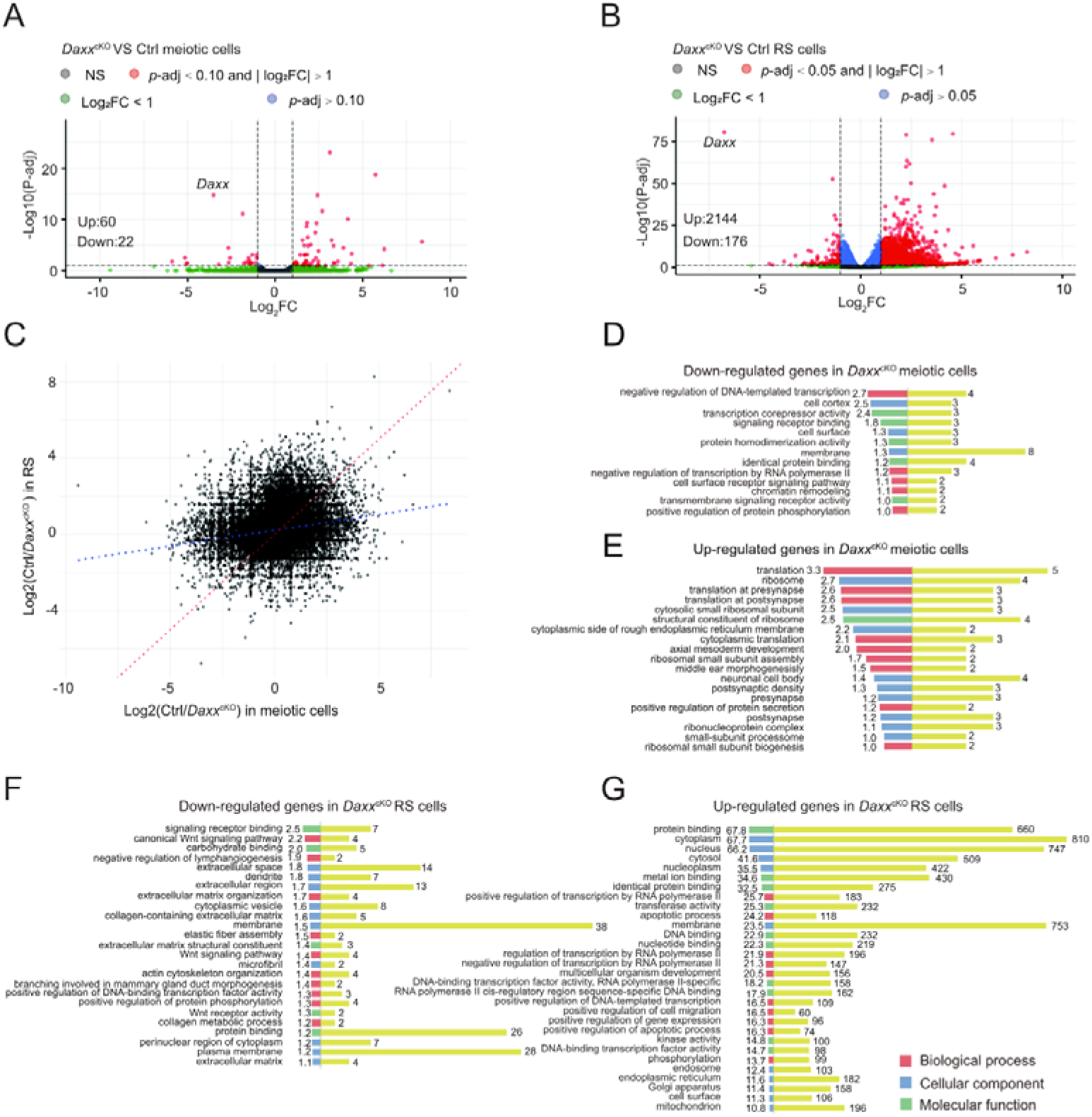
*Daxx* deletion altered the transcriptomes of meiotic cells and round spermatids. (A) Comparative transcriptomic profiling via scatter plot visualization revealed gene expression dysregulation in *Daxx*^cKO^ meiotic cells when compared with the controls. The original data are available in Table S2. (B) Comparative transcriptomic profiling via scatter plot visualization revealed gene expression dysregulation in *Daxx*^cKO^ RS when compared with the controls. The original data are available in Table S2. (C) Scatter plot of all pairwise log2-fold change in gene expression in the absence of DAXX between meiotic cells and RS. The red dashed line is the Y = X axis and the blue dashed line is the regression line. (D) Gene ontology (GO) enrichment analysis of the down-regulated genes of meiotic cells in the absence of DAXX. The numbers on the right represent the number of genes in the pathway and the numbers on the left represent -log10 (*P*-value). (E) GO enrichment analysis of the up-regulated genes of meiotic cells in the absence of DAXX. The numbers on the right represent the number of genes in the pathway and the numbers on the left represent -log10 (*P*-value). (F) GO enrichment analysis of the down-regulated genes in RS in the absence of DAXX. The numbers on the right represent the number of genes in the pathway and the numbers on the left represent -log10 (*P*-value). (G) GO analysis enrichment of the up-regulated genes of RS in the absence of DAXX. The numbers on the right represent the number of genes in the pathway and the numbers on the right represent -log10 (*P*-value).

### Transposons are derepressed in *Daxx*^cKO^ mice

Besides transcriptional repression, DAXX also plays a critical role in silencing ERVs (endogenous retroviral elements) in murine embryonic stem cells ^42–45^. Other studies have also demonstrated that the loss of DAXX in the mouse pancreas has been linked to the derepression of ERVs, leading to increased expression of nearby protein-coding genes ^46^.

To investigate the effects of DAXX deletion on TEs expression, we conducted transcriptomic analysis and found significant upregulation of two retrotransposon subsets, that 27 differentially expressed ERVs (20 ERVK, 3 ERVL and 4 ERV1) and 12 differentially expressed LINE1 in *Daxx*^cKO^ meiotic cells. A different pattern emerged in *Daxx*^cKO^ post-meiotic cells, that 34 differentially expressed ERVs (25 ERVK, 4 ERVL and 5 ERV1) and 8 differentially expressed LINE1 showed a significant increase with a *p*-value < 0.05 and log2FC > 1 (Figure. 6A). Evolutionarily young LINE1s are capable of autonomous transposition and their activation compromises genomic stability. In *Daxx*^cKO^ meiotic cells, young LINEs, including L1Md_A, L1Md_Gf and L1Md_Tf, were upregulated, however, they were not upregulated in *Daxx*^cKO^ RS (Figure. 6A). We further examined the expression of LINE1 protein in *Daxx*^cKO^ testes and found that LINE1 ORF1P protein was barely detectable in control tubules as expected. In contrast, *Daxx*^cKO^ testes exhibited a much higher abundance of LINE1 ORF1P in spermatocytes (Figure. 6C and 6D). Correspondingly, western blot analysis reported an elevated LINE1 ORF1P protein in *Daxx*^cKO^ testes (Figure. 6E and 6F). Overactivated LINE1 ORF1P was observed in germ cells of *Daxx*^cKO^ testes as early as on P14D when late zygotene or early pachytene cells appeared during the first wave of spermatogenesis (Figure. 6G). DAXX deficiency also led to overexpression one specific ERV subfamily IAPEy in *Daxx*^cKO^ meiotic cells (which phenocopied the *Miwi2* ^17^- and *Spocd1* ^16^ - deficient spermatocytes) as well as *Daxx*^cKO^ post-meiotic cells (Figure. 6A). MMERVK10c was also significantly upregulated in *Daxx*^cKO^ RS (Figure. 6A). In conclusion, DAXX deficiency results in derepression of a large number of TE families, particularly the young LINEs in meiotic cells.

**Figure 6.**
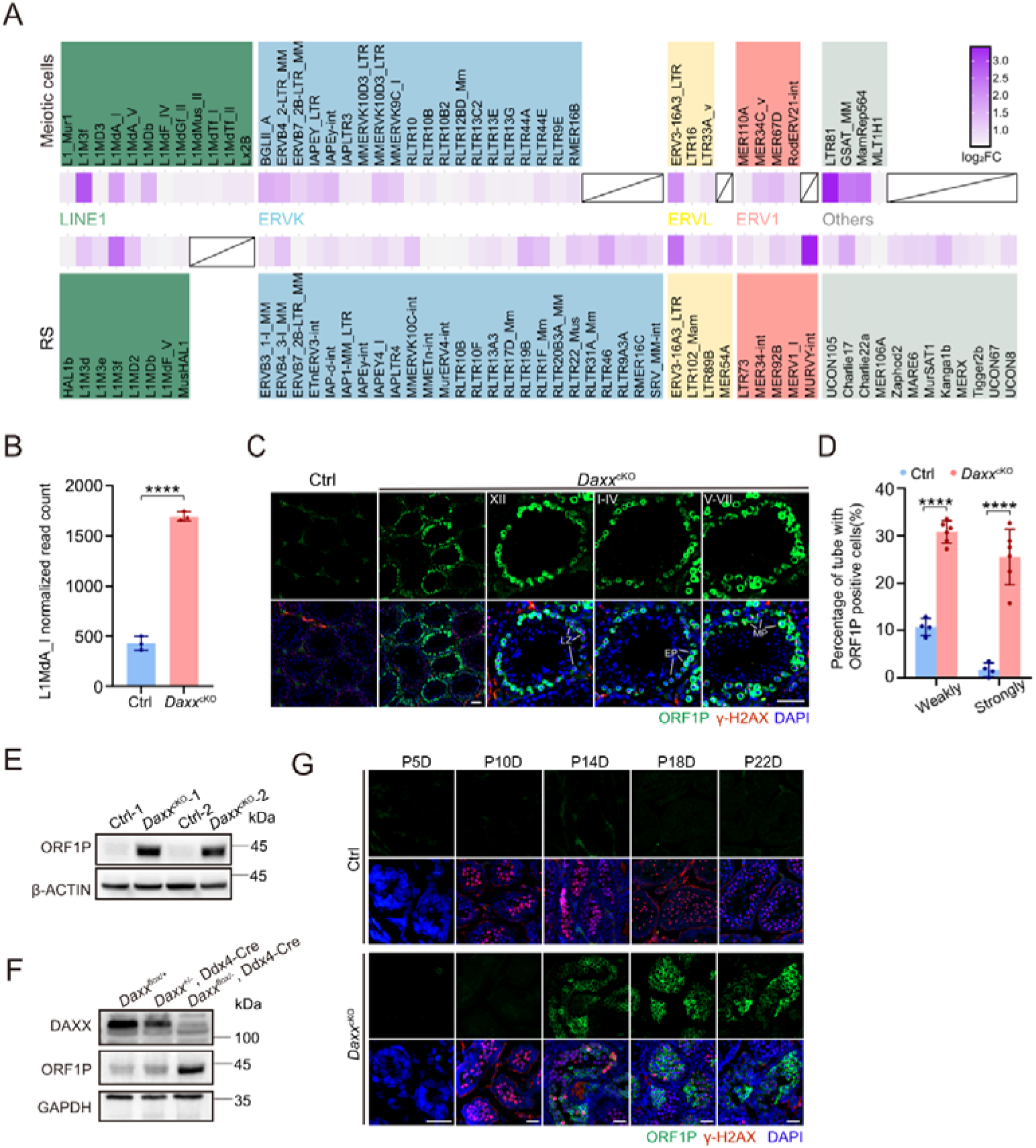
Young LINE1 were over-activated in meiotic cells of *Daxx*^cKO^ males. (A) RNA-seq data revealed transposable elements are upregulated in *Daxx*^cKO^ meiotic cells and RS, with heatmaps depicting fold-change of expression. *p*-value < 0.05 and log_2_FC > 1 are shown. The original data are available in Table S3. (B) Normalized read counts for L1MdA_I from RNA-seq data in meiotic cells from control and *Daxx*^cKO^. *\*\*\*\*p* = 8.02×10^-15^ by Benjamini–Hochberg adjusted Wald test and the original data are available in Table S3. (C) Representative images of immunofluorescence staining of ORF1P (green), γH2AX (red), and DAPI (blue) on testicular sections from 4-mounth-old control and *Daxx*^cKO^ mice. The co-stained γH2AX was used for staging of spermatocytes. LZ: late-zygotene; EP: early-pachytene; MP: middle-pachytene. Scale bar, 50 μm. (D) Quantification of ORF1P positive seminiferous tubules. *\*\*\*\*p* < 0.0001 by two-tailed Student’s *t*-test. (E) Western blot analyses of ORF1P protein in control and *Daxx*^cKO^ testes from adult mice. β-ACTIN serves as the loading control. (F) Western blot analyses of ORF1P protein in WT, heterozygote and homozygote testes from adult mice. GAPDH serves as the loading control. (G) Representative images of immunofluorescence staining of ORF1P (green), γH2AX (red), and DAPI (blue) on testicular sections at P10D, P14D, P18D and P22D from control and *Daxx*^cKO^ mice. The co-stained γH2AX was used for staging of spermatocytes. Scale bar, 50 μm

### DAXX is required for maintaining LINE1 DNA methylation but not piRNA pathway

The overactivation of retrotransposons in *Daxx*^cKO^ mice closely resembles the phenotype observed in piRNA pathway-deficient mutants ^47–49^. It is well documented that pachytene-stage piRNAs derive from a defined set of 100 TE-poor genomic regions defined as piRNA clusters ^50^ and its function are to silence active transposons by regulating the translation of their target transcripts in the spermatocytes ^51^. Therefore, we next asked whether DAXX functions in repressing transposons via the piRNA pathway. We examined the pachytene piRNA-related proteins MIWI (PIWIL1) and MILI (PIWIL2) by western blot and IF, and found that both of them exhibited similar expression in *Daxx*^cKO^ testes to the controls (Figure. S7).

To clarify whether piRNA biogenesis is intact after DAXX deletion, we performed small RNA sequencing on meiotic cells and RS from control and *Daxx*^cKO^ testes and found that piRNAs were present in *Daxx*^cKO^ testes with comparable length distribution between control and *Daxx*^cKO^ testes (Figure. 7A). We also didn’t find impacts of *Daxx*-deficiency on mapped piRNAs (Figure. 7B, S8A and S9A), either their counts or amplification (Figure. S8 and S9). It’s noteworthy that a significant increase of L1MdA-derived antisense piRNAs was detected in *Daxx*^cKO^ meiotic spermatocytes (Figure. S8A), which may be caused by the more abundant L1MdA transcripts in *Daxx*^cKO^ meiotic spermatocytes. These findings suggest that DAXX either functions independently of piRNA biogenesis or serves as a nuclear execution of piRNA-guided silencing TEs during spermatogenesis.

**Figure 7.**
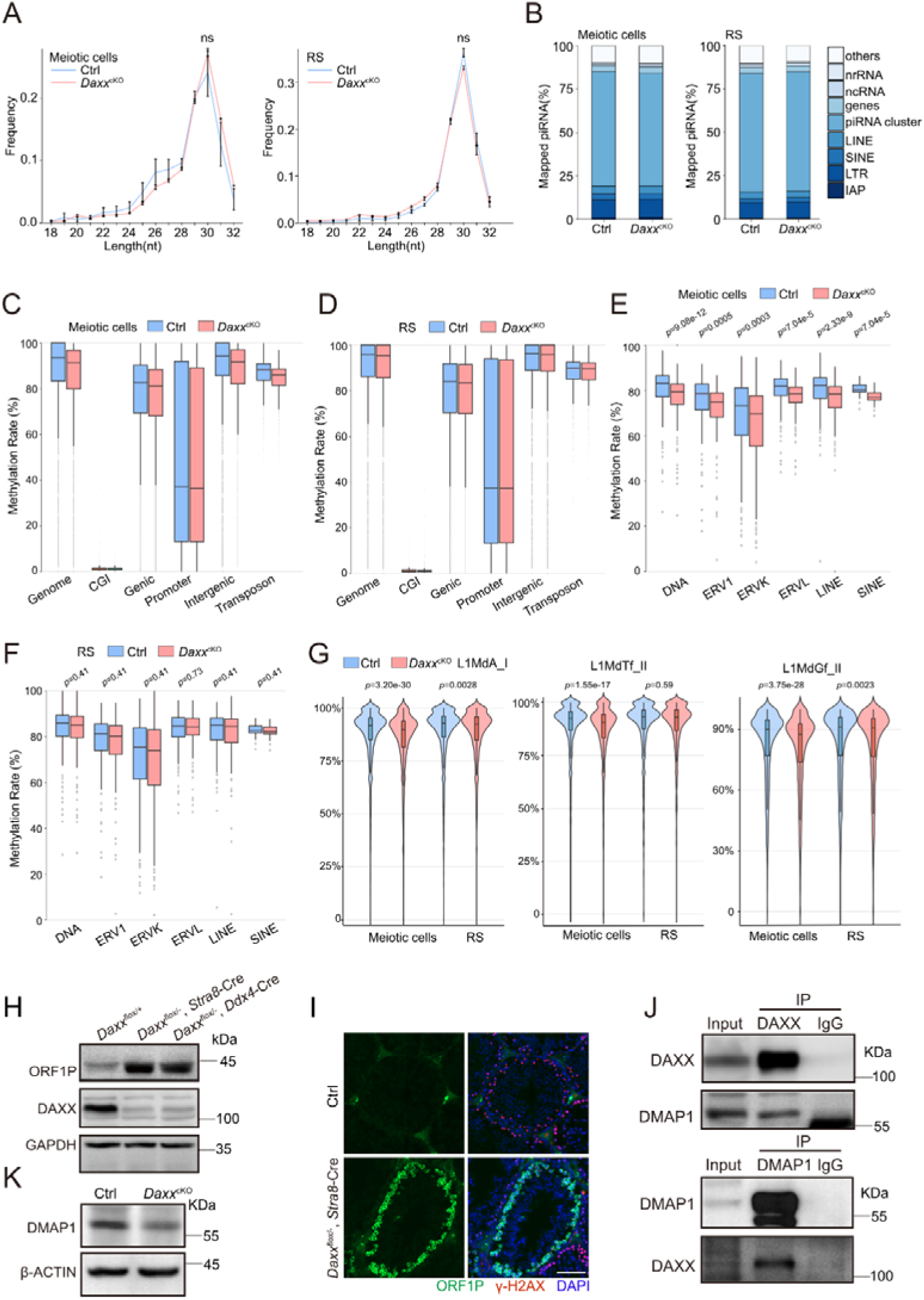
DAXX is required for DNA methylation maintenance of young LINE1 but not piRNA pathway. (A) Length (nt) distribution of small RNAs from purified meiotic spermatocytes and RS of control and *Daxx*^cKO^ mice. Data represent the mean and *s.e.m.* No significant differences were observed (Bonferroni adjusted two-tailed Student’s *t*-test). The original data are available in Table S4. (B) Genomic annotation of total piRNA from purified meiotic spermatocytes and RS from control and *Daxx*^cKO^ mice. The original data are available in Table S4. (C, E) Percentages of CpG methylation levels of the indicated genomic compartments in meiotic cells (C) and RS (E) from control versus *Daxx*^cKO^. The original data are available in Table S5. (D, F) Percentages of CpG methylation levels of the indicated classes of transposable elements in meiotic cells (D) and RS (F) from control versus *Daxx*^cKO^. The original data are available in Table S5. (G) Percentages of CpG methylation levels of young LINE1 elements in meiotic cells and RS from control versus *Daxx*^cKO^. The original data are available in Table S5. (H) Western blot analyses of ORF1P protein in control, *Daxx*^flox/-^, *Stra8*-Cre testes and *Daxx*^flox/-^, *Ddx4*-Cre from adult mice. GAPDH serves as the loading control. (I) Representative images of immunofluorescence staining of ORF1P (green), γH2AX (red), and DAPI (blue) on testicular sections from adult control and *Daxx*^flox/-^, *Stra8*-Cre mice. Scale bar, 50 μm. (J) Co-IP analysis of DAXX with DMAP1 from testicular protein extracts. (K) Western blot analyses of DMAP1 protein in control and *Daxx*^cKO^ testes from adult mice. β-ACTIN serves as the loading control. Data in C-G, Statistics: Benjamini–Hochberg adjusted Wilcoxon test.

DNA methylation serves as the principal epigenetic modification responsible for repression of TEs. To interrogate DAXX’s regulatory influence on TEs DNA methylation, whole-genome bisulfite sequencing (WGBS) was implemented on isolated meiotic cells and RS from control and *Daxx*^cKO^ testes. Indeed, we observed no significant change in global methylation in *Daxx*^cKO^ RS, but a statistically significant yet modest decrease in methylation was observed in genome genic, intragenic, promoter and transposon regions of *Daxx*^cKO^ meiotic cells (Figure. 7C and 7D). DNA methylation patterns was also assessed at individual TE families. Globally, we observed a significant hypomethylation in LINE, ERVK, ERVL, ERV1, and SINE of *Daxx*^cKO^ meiotic cells, but there was no significant change in *Daxx*^cKO^ RS (Figure. 7E and 7F). Consistently, young LINE1 subfamilies L1Md_A, L1Md_Tf and L1Md_Gf displayed a significant demethylation in *Daxx*^cKO^ meiotic cells but not in *Daxx*^cKO^ RS (Figure. 7G).

Notably, LINE1 are also upregulated in spermatocytes of *Daxx*^flox/-^, *Stra8*-Cre mice (Figure. 7H and 7I), demonstrating that their activation is not attributable to defects in de novo DNA methylation. We previously hypothesized that the LINE-1 activation and hypomethylation in meiotic cells of *Daxx*^cKO^ mice might arise from defects in de novo methylation in PGCs rather than impaired maintenance of methylation during meiosis. To test this, we analyzed LINE1 expression in *Stra8*-cre mediated *Daxx* deletion testis (*Stra8* expression initiates at P5D in male mice, whereas genome-wide de novo methylation is largely complete at P2.5D ^15^). The observed upregulation of LINE1 in *Daxx*^flox/-^, *Stra8*-Cre supports that the hypomethylation is indeed attributable to maintenance defects in meiotic cells.

Biochemical assays demonstrate that DMAP1 (DNMT1-associated protein 1) strongly enhances the methyltransferase activity of DNMT1, implying its role as a cofactor that facilitates methylation ^52^. In vitro biochemical characterization revealed a direct interaction between DMAP1 and DAXX ^53^, suggesting potential functional collaboration. We confirmed the co-precipitation of DAXX with DMAP1 in vivo. DMAP1 was identified in DAXX-immunoprecipitated (IP) complexes from testis lysates, and their association was confirmed by reciprocal IP experiments (Figure. 7J). In addition, western blot analysis of the DMAP1 protein was decreased in *Daxx*^cKO^ testes (Figure. 7K).

## DISCUSSION

DAXX serves as an essential protein with a multifaceted interactome, whose disruption results in embryonic lethality at E6.5 accompanied by extensive apoptosis, demonstrating its indispensable role in embryonic development ^24^. In the current work, we have carried out an in-depth study on the role of DAXX in mouse spermatogenesis. Germ cell-specific *Daxx* deletion models coupled with multi-dimensional profiling has allowed us to illustrate DAXX as a vital factor in the mammalian spermatogenesis (Figure. 8).

**Figure 8.**
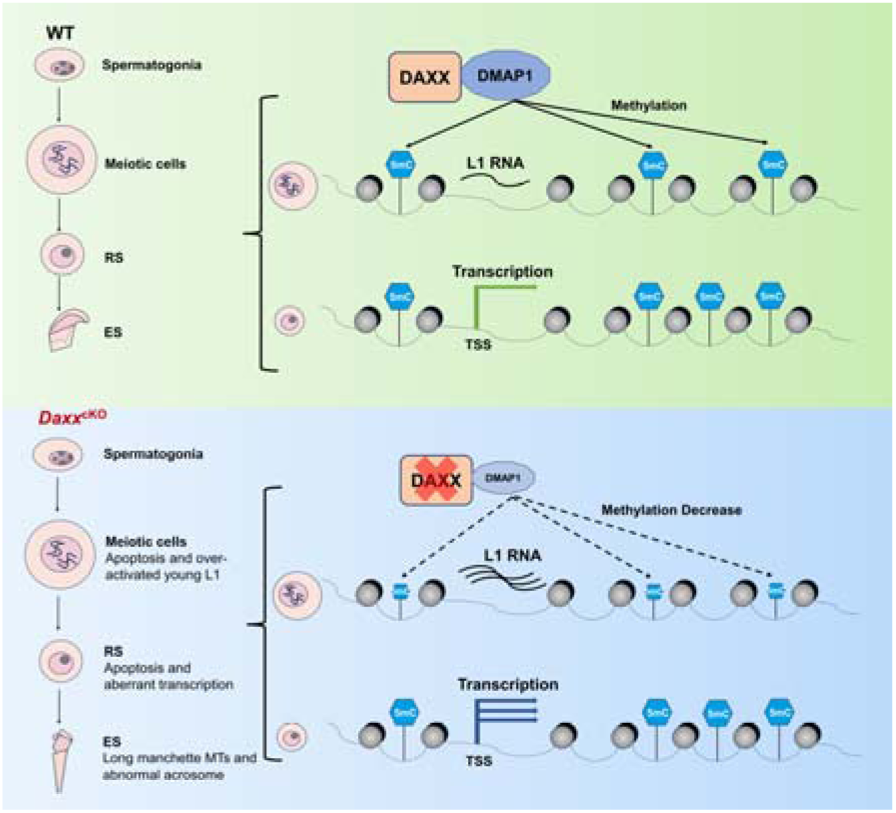
**Working model of the role of DAXX in spermatogenesis.** DAXX interacts with DMAP1 to maintain DNA methylation of LINE1 in meiotic cells.

The *Daxx*^cKO^ male mice exhibited germ cells loss and an age-dependent decline in fertility. The morphology of the spermatozoa was highly altered and the sperm concentration was strongly decreased in *Daxx*^cKO^ mice. In particular, excessive elongation of the manchette MTs of spermatids occurs in *Daxx*^cKO^ mice. The manchette assembles by microtubule-actin to form a dynamic cytoskeleton, which creates a transient intracellular structure during the phase of sperm head shaping ^54–56^. Manchette assembly initiates at step 8 of spermiogenesis, temporally coupled with nuclear elongation, and undergoes disassembly during step 13 to 14 concurrent with nuclear architectural finalization. Aberrant manchette structure impair nuclear shaping, resulting in defective sperm head morphogenesis ^37,55^. We found DAXX plays a role in the organization of manchette MTs network and a single cluster consisting of significantly down-regulated actin cytoskeleton organization in *Daxx*^cKO^ RS. However, the in-depth reasons for the defect deserve continued investigation.

Numerous articles reported the association of DAXX with ERVs, but few have reported the correlation between DAXX and young LINE1. Here we have analyzed the expression of the repetitive elements during spermatogenesis. Depletion of DAXX triggered mass retrotransposons transcriptional activation in both meiotic cells and RS, therein, the dysregulation of MMERVK10C-int ^9^ and L1MdA ^5^ retrotransposons are likely implicated in infertility. Young LINE1 families exemplified by L1Md_A, L1Md_Tf and L1Md_Gf were up-regulated in meiotic cells, however, they were not up-regulated in *Daxx*^cKO^ post-meiotic cells. Considering that DAXX deficiency does not significantly alter DNA methylation patterns at young LINE1 in post-meiotic cells, we hypothesized that some meiotic cells escaped transposon derepression in *Daxx*^cKO^ spermatocytes and passed through meiosis into RS. These demonstrate that DAXX’s transcriptional repression of young LINE1 during meiosis is not globally required.

*Daxx* deletion triggers LINE1 activation and apoptosis in germ cells, and a subsequent decrease in post-meiotic cell populations, yet conventional cytological assessment fails to detect obvious disruption of chromosomal behaviors during meiosis, suggesting that defects likely arise after diplotene stage in the progression to RS or subtler functional impairments.

Our study established that DAXX interacts with DMAP1 in mouse testes, and specifically demonstrates that DAXX knockout significantly reduces DMAP1 protein levels in testicular lysates. This observation validates the proposition proposed by Ryuta Muromoto et al., wherein DAXX protects DMAP1 from degradation in vivo. DNMT1, an essential enzyme for DNA methylation maintenance, exhibits ∼2.5-fold elevated methyltransferase activity upon DMAP1 binding in vitro. We therefore proposed that during meiosis, DAXX stabilizes DMAP1, thereby enabling DMAP1-dependent potentiation of DNMT1 activity. This pathway maintains DNA methylation patterns, consequently ensuring effective suppression of young LINE1 activation. The limitation lies in the unresolved chromosomal localization of DAXX and DMAP1. While immunofluorescence staining has delineated DAXX’s subcellular distribution at a coarse resolution, no effective antibody for immunoprecipitation precluded precise mapping. To address this, future work will be needed to employ tagged mouse models coupled with IP-MS and ChIP-seq to systematically investigate the correlation of DAXX/DMAP1 binding and DNA hypomethylation on LINE1.

To delineate *Daxx*-dependent chromatin remodeling effects, ATAC-seq of meiotic spermatocytes and RS revealed accessible chromatin domains, with no significant in peaks distribution across genomic regions between the two genotypes of males (Figure. S10A). Comparative chromatin accessibility analysis between control and *Daxx*^cKO^ meiotic cells revealed 4,473 differentially accessible regions, comprising 330 regions open and 4,143 closed. We next evaluated all differentially accessible regions and found that 50.91% (168 of 330) of the differentially open regions contained TEs sequence, 15.21% (630 of 4,143) of the differentially closed regions contained TEs sequence. Notably, young LINE1 were preferentially localized to differentially open regions (21 regions) over differentially closed regions (9 regions) (Figure. S10B). The above findings suggest that DAXX deletion may activate the expression of young LINE1 elements by altering chromatin accessibility in spermatocytes.

Considering the specific localization of DAXX in the centromere regions of the spermatocytes, we speculate that DAXX is likely to be involved in the first meiotic division as it has been demonstrated that ATRX recruits DAXX to pericentric heterochromatin in preovulatory oocytes ^57^ and loss of maternal ATRX function at the metaphase II stage disrupts chromosome alignment and adversely affects female fertility ^57–59^. Interestingly, oocytes from *Daxx*^cKO^ females retain the ability to support preimplantation embryonic development (Figure. S11A-11F). The same as this, DAXX deficiency does not affect chromosome arrangement in meiotic metaphase I cells of male mice (Figure S11G). This reveal that DAXX and ATRX exhibit distinct functional contributions to oocyte development.

## METHOD DETAILS

### Mice

Mice were maintained and housed at the standard animal facility of Wuhan University according to the Regulations for the Administration of Affairs Concerning Experimental Animals of China. All animal experiments strictly adhered to the ethical standards and protocols approved by The Laboratory Animal Care and Use Committee of the Wuhan University. *Daxx*^flox/flox^ mice were generated by CRISPR-Cas9 gene-editing approach. Testis-specific *Daxx* depletion was achieved by crossing *Daxx*^flox/flox^ mice with *Ddx4*-Cre (Jackson Laboratory 006954) mice. The *Stra8*-GFPCre mouse was gifted Dr. Ming-Han Tong ^60^.

### Production of the DAXX antibody

For polyclonal antibody production, a recombinant mouse *Daxx* antigen was engineered by cloning the corresponding cDNA in-frame into the Pet42b expression vector, generating an N-terminal amino acids 626 to739 fused with an 6x His tag. Following IPTG induction in BL21-CodonPlus (DE3) E. coli, the His-tagged protein was purified via nitrilotriacetic acid affinity chromatography. After extensive dialysis against PBS, the antigenic preparation was utilized for rabbit immunization. The resulting antisera demonstrated specific immunoreactivity, and used for western blotting, immunofluorescence staining.

### Histological processing and analysis

Tissues for each genotype were removed immediately after euthanasia, fixed in specialized fixative for 24 hours, and embedded in paraffin after dehydration. Sections of 5 μm thickness were microtome-prepared and mounted on slides. Following xylene-mediated deparaffinization, slides were stained with hematoxylin and eosin for histological analysis.

### Chromosome spread and immunofluorescence staining

Spermatocyte spreads were prepared as previously described ^61^. Primary antibodies used for chromosome spread immunofluorescence were as follows: rabbit anti-DAXX (1:100 dilution), mouse anti-SYCP3 (1:200 dilution) and anti-autoantibody against centromere (1:200 dilution). Primary antibodies were detected with Alexa Fluor 488-, 594-, or 405 conjugated secondary antibodies for 1 hour at room temperature. The slides were washed with TBS four times and mounted with DAPI.

### Immunofluorescence staining and TUNEL assay

Samples of mouse testis were fixed in 4% phosphate buffered PFA overnight. The fixed samples were dehydrated, embedded into Optimal Cutting Temperature compound (OCT) and cut into 5-8 μm sections on a cryostat. Primary antibodies used for testicular sections immunofluorescence were as follows: mouse anti-γH2AX (1:300 dilution), LINE1 ORF1 (1:200 dilution), PIWIL1/MIWI (1:200 dilution), PIWIL2/MILI (1:200 dilution), YBX2 (1:300 dilution), Histone H3 (1:200 dilution), Protamine P1 (1:100 dilution), TNP1 (1:100 dilution), α-TUBLIN (1:100 dilution), PLZF (1:300 dilution), SOX9 (1:200 dilution), TRA98 (1:100 dilution). TUNEL assay was performed on testicular sections using the cell death detection kit according to the manufacturer’s instructions. Primary antibodies were detected with Alexa Fluor 488-, or 594-conjugated secondary antibodies for 1 hour at room temperature. The slides were washed three times with TBS and coverslipped with DAPI-supplemented antifade mounting solution. Structured illumination microscopy (SIM) was performed using a ZEISS Axio Observer 3 microscope. Images were then processed using ZEN 2.6 lite for analysis.

### Western blot

Freshly excised tissues from C57BL/6 mice underwent mechanical disruption in ice-cold lysis buffer [62.5 mM Tris-HCL (pH 6.8), 3% SDS, 10% glycerol, and 20% β-mercaptoethanol] supplemented with protease inhibitor cocktail. After ultrasound on the ice, the tissue extracts were centrifuged at 20,000 g for 10 min at 4°C. The supernatant extracts were combined with protein loading buffer and denatured at 100 °C for 10 minutes. Equal amounts of protein lysates were resolved under denaturing conditions via SDS-polyacrylamide gels, followed by the bands to polyvinylidene fluoride (PVDF) membranes (0.45 μm pore size). Immunoreactive PVDF membranes were detected and analyzed with Super Signal™ West Pico PLUS Chemiluminescent Substrate and G: Box Chemi-XRQ GENE Sys. Quantitative analysis of target proteins was calibrated against β-ACTIN/GAPDH.

### Isolation of mouse germ cells

Spermatocytes were isolated by using the fluorescence-activated cell sorting method ^29,30^. Briefly, the tunica albuginea of testes was removed in DMEM, and the testes were digested with collagenase type I (500 μg/ml, 32 °C, 15-20 min, orbital shaking at 100pm) then further digested with trypsin (200 μg/ml) and DNase I (20 μg/ml) at 32 °C for 5 to10 min, and the digestion was terminated with 10% fetal bovine serum. The cell suspension underwent the 70um cellular filter, and low-speed centrifugation (500 g, 4°C, 5 min) for pellet isolation. The cells were resuspended in DMEM with 10% fetal bovine serum, fluorescently labeled with Hoechst 33342 and sorted via BD FACS Aria II using 355 nm excitation and 450 nm emission parameters.

### Computer-assisted motility analysis (CASA)

One epididymis was collected from each adult mouse and placed in a 1.5 ml Eppendorf tube. The tissue was minced and incubated with 1 ml of pre-warmed 1x PBS at 37°C for 30 minutes to allow sperm to swim out. Subsequently, sperm counts were determined using a hemocytometer. Quantitative kinematic profiling of spermatozoa was conducted using the motility module of a clinically validated multiparametric andrological assessment system (BEION platform, version 4.20).

### mRNA sequencing and analysis

We collected meiosis cells and RS from adult mice via FACS as described above. RNA was extracted according to the instruction manual of the TRlzol Reagent. All RNA libraries were constructed at the same time using the NEBNext Ultra Directional RNA Library Prep Kit for Illumina.

For analysis of differentially expressed genes, Adapters and low qualities bases were trimmed with Trim Galore ^62^ version 4.9, parameters:–paired–length 25--stringency 4--clip_R1 12 --clip_R2 12). Then, clean sequence reads were mapped to the UCSC mm39 genome with default parameters using HISAT2(2.2.1) ^63^, mapped reads per gene were counted with FeatureCount (1.5.3) ^64^, providing a GTF file acquired from UCSC RefSeq annotation, and differential expression analysis was performed by DESeq2(1.46.0) ^65^ with R (4.4.1). The analysis of Gene ontology showing differential expression genes was carried out using DAVID 6.8. For analysis of differentially expressed Transposable element, similar to the analysis mentioned above, and the GTF file was acquired from Gale-Hammell lab (https://labshare.cshl.edu/shares/mhammelllab/www-data/TEtranscripts/TE_GTF/mm39_rmsk_TE.gtf.gz).

### Small RNA sequencing and analysis

The tool cutadapt (4.9) was employed to trim the adaptor sequences (AGATCGGAAGAGCACACGTCT) from the 3′ ends of the raw fastq files ^62^ with parameters(-m 16 --trimmed-only), Small RNA reads of 18–32 nt for each sample were retrieved as described for piRNA analyses ^16^. These reads were aligned to miRNA retrieved from miRBase, non-coding RNAs, piRNA clusters and genomic transposable element sequences retrieved from RepeatMasker (mm39) ^50^. When mapping reads to retrotransposons elements, up to three mismatches were permitted. Reads of piRNA that were 25–32 nt were classified based on annotations, and those not aligning with any known genomic elements were grouped as ‘other’. For the analysis of individual piRNA expressions, only mapped reads between 25–32 nt and exceeding 10 counts were considered and visualized in scatter plot. For Ping-pong analysis of piRNAs, the consensus sequences of L1MdTf_I, L1MdGf_I, L1MdF_I, L1MdA_I, IAPEY and IAPEZI were retrieved from Repbase ^66^, and mapping the FASTA format sequences to the above consensus sequence, allowing up to three mismatches as described in. Then, calculate the distances of 5’ end of sense/antisense piRNAs and count first and 10^th^ nucleotide from 5’ end of piRNA.

### DNA methylation analysis

Adapters and low qualities bases were trimmed with Trim Galore (parameters: --paired --length 25 --clip_R2 8), the clean reads were aligned to the UCSC mm39 genome carried out with Bismark (v0.24.2) ^67^ with the following set of parameters: –score_min L,0,-0.4, which incorporates Bowtie2 (2.5.4) ^68^. CpG methylation calls were extracted from the deduplicated mapping output using the Bismark methylation extractor. Only CpGs covered by a minimum of three reads were retained for downstream analyses in each replicate.

The methylation of genomic features was calculated by intersecting CpG positions with genomic annotations using bedtools (v2.31.1) ^69^ intersect, and averaging methylation of each feature. We used the CpG-island annotation, RefSeq gene annotation, and Transposable element annotation mentioned above, promoters were defined as -2kb around RefSeq TSS, and we defined intergenic regions as genomic regions that do not overlap with any of the previous annotations. For the analysis of methylation levels of different repClass as well as individual transposons, we processed the aforementioned GTF file and overlaped with CpG positions useing bedtools intersect, and Python (3.12.7) was used to calculate the average methylation rate. The results were then visualized in RStudio. The methylation conversion rate was calculated by mapping all reads to the spiked-in CpG unmethylated lambda.

### Assay for transposase-accessible chromatin and analysis

Meiotic spermatocytes and RS were sorted via FACS as described above. Isolated cells underwent PBS washes followed by nuclear prepared through lysis buffer [10 mM Tris-HCl (pH 7.4), 10 mM NaCl, 3 mM MgClLJ, 0.1% NP-40] with 10 min ice incubation and centrifugation (500 g, 5 min, 4°C). Nuclei pellets were resuspended in Tn5 transposase reaction mix for 37°C tagmentation (10 min). DNA fragments were then immediately purified with ATAC DNA extract beads prior to PCR amplification with dual-indexed primers using TruePrep DNA Library Prep Kit V2 for library construction.

Quality assessment and adaptor sequences removement of raw data were using Trim Galore (parameters: -- paired --length 25 --clip_R2 8) reads were aligned to the UCSC mm39 genome using Bowtie2 (2.5.4) ^68^. Samtools (version 1.8) was employed for sorting, format conversion, and filtering of alignment reads. Chromosomal reads from the Mitochondria were excluded ^70^, and duplicates were removed using Sambamba (version 1.0.1) ^71^. Reads aligned to multiple locations were discarded, preserving only those uniquely aligned to chromosomal locations and properly paired. Peak calling was performed using MACS2 (v2.2.7.1) ^72^, and annotation was carried out using HOMER (5.1) ^73^. To identify differentially chromatin regions, we used the R package DiffBind (3.16.0), and performed an overlap analysis with transposon regions to examine the distribution of these regions.

### Statistical analysis

Quantitative data processing and graphical visualization were executed utilizing GraphPad Prism version 9.51. No samples or animals were excluded from the final dataset, and sample size calculation was not conducted prior to experimentation. Data are shown as mean ± SD unless noted. The statistical methods and p-values used in this study are described in corresponding figure legends or results.

## DATA AVAILABILITY

The data that support the findings of this study are available in NCBI at https://www.ncbi.nlm.nih.gov/geo/query/acc.cgi?acc=GSE293055. This study did not generate any custom code. Any additional information required to reanalyze the data reported in this paper is available from the lead contact upon request. All data necessary to replicate the findings of this study are available from the lead contact upon request.

## ACKNOWLEDGMENTS

This research was supported by the National Natural Science Foundation of China (32270905), Natural Science Foundation of Hubei Province - Innovation Group project (2024AFA018), Shenzhen Municipal Science and Technology Innovation Committee (JCYJ20240813111406009), Zhongnan Hospital of Wuhan University (ZNLH202206) and Yichun Maternal and Child Health Hospital to Mengcheng Luo, and by the Ministry of Science and Technology of China (2023YFA0913400), the National Natural Science Foundation of China (32230020 and 31991161), Basic Research Program Based on Major Scientific Infrastructures, CAS (JZHKYPT-2021-05) and the New Cornerstone Science Foundation to Guohong Li. The work was also supported by the Wuhan University open subsidies for large instruments and equipment to Rong Liu. We thank Qian Liu from the Animal Experimental Center, College of Life Sciences, Wuhan University for assistance with animal experiments. We also thank professor Ming-Han Tong from Chinese Academy of Sciences for kindly providing the *Stra8*-GFPCre mouse.

## AUTHOR CONTRIBUTIONS

Conceptualization, L.M., L.G., L.R. and T.Y; methodology, L.Z., X.C., S.J.; Investigation, L.Z., X.C., S.J., T.C., M.D., C.J., S.Q., G.W., Y.W., H.M. and H.L.; writing – original draft, L.Z.; writing – review & editing: L.R.; computational analyses, S.J.; funding acquisition, L.M., L.G. and L.R.; resources: L.M., L.G. and L.R.; supervision, L.M., L.G., L.R. and T.Y.

## SUPPLEMENTAL INFORMATION

Document S1. Figures S1–S11

Table S1. List of primers used in this study

Table S2. List of differentially expressed genes

Table S3. List of differentially expressed transposons

Table S4. piRNA analysis

Table S5. Whole-genome methylation analysis

Table S6. ATAC-seq analysis

Table S7. List of reagent or resource

## Supplemental information

**Figure S1.**
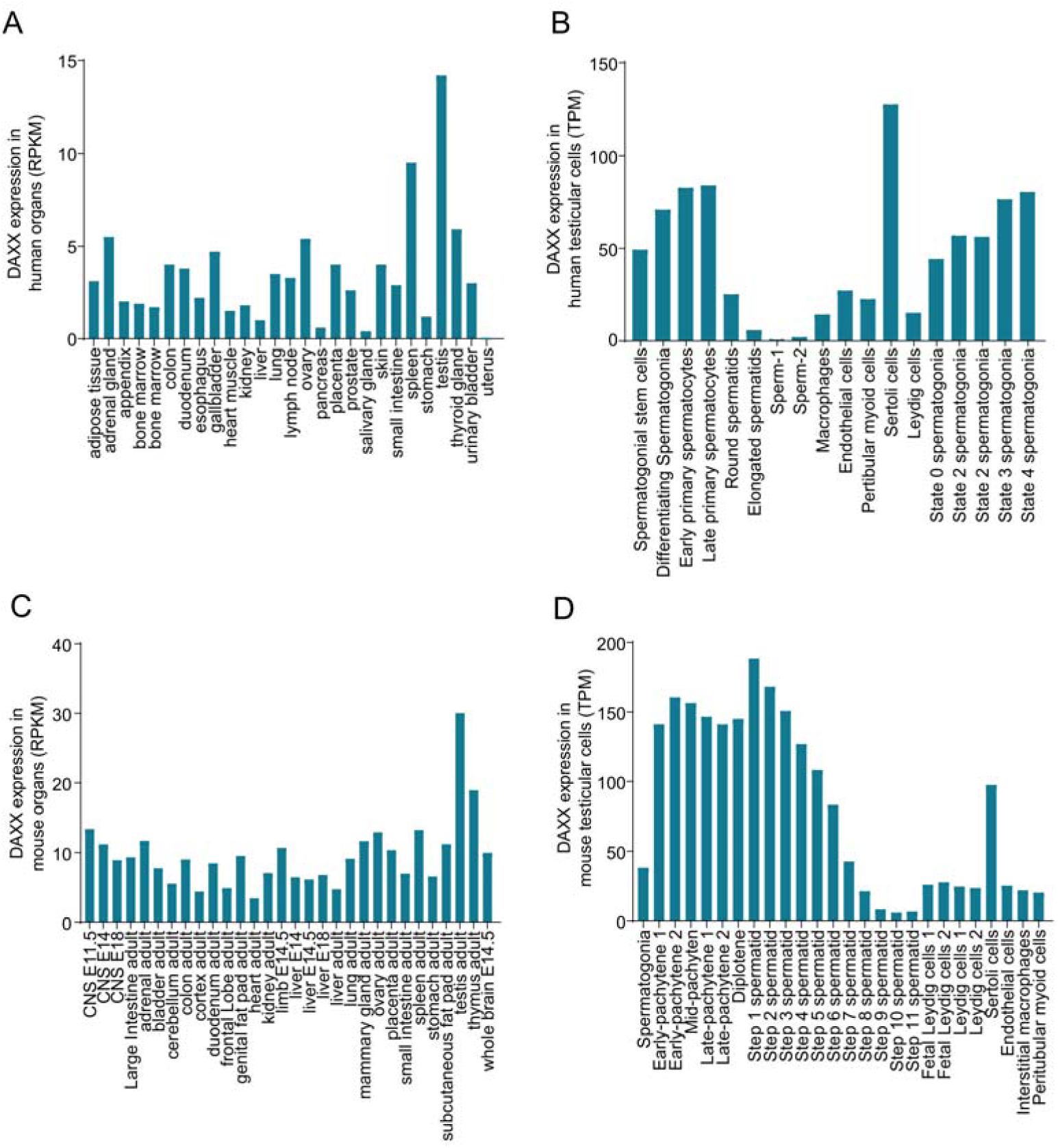
DAXX transcript are highly expressed in human and mouse testes. Related to Figure 1. (A) The *Daxx* expression profile among various human organs/tissues. RPKM: Reads Per Kilobase per Million mapped reads. (B) The *Daxx* expression profile among the human testicular cells based on published single-cell RNA sequencing data. TPM: Transcripts Per Million. (C) The *Daxx* expression profile among various mouse organs/tissues. RPKM: Reads Per Kilobase per Million mapped reads. (D) The *Daxx* expression profile among the mouse testicular cells based on published single-cell RNA sequencing data. TPM: Transcripts Per Million.

**Figure S2.**
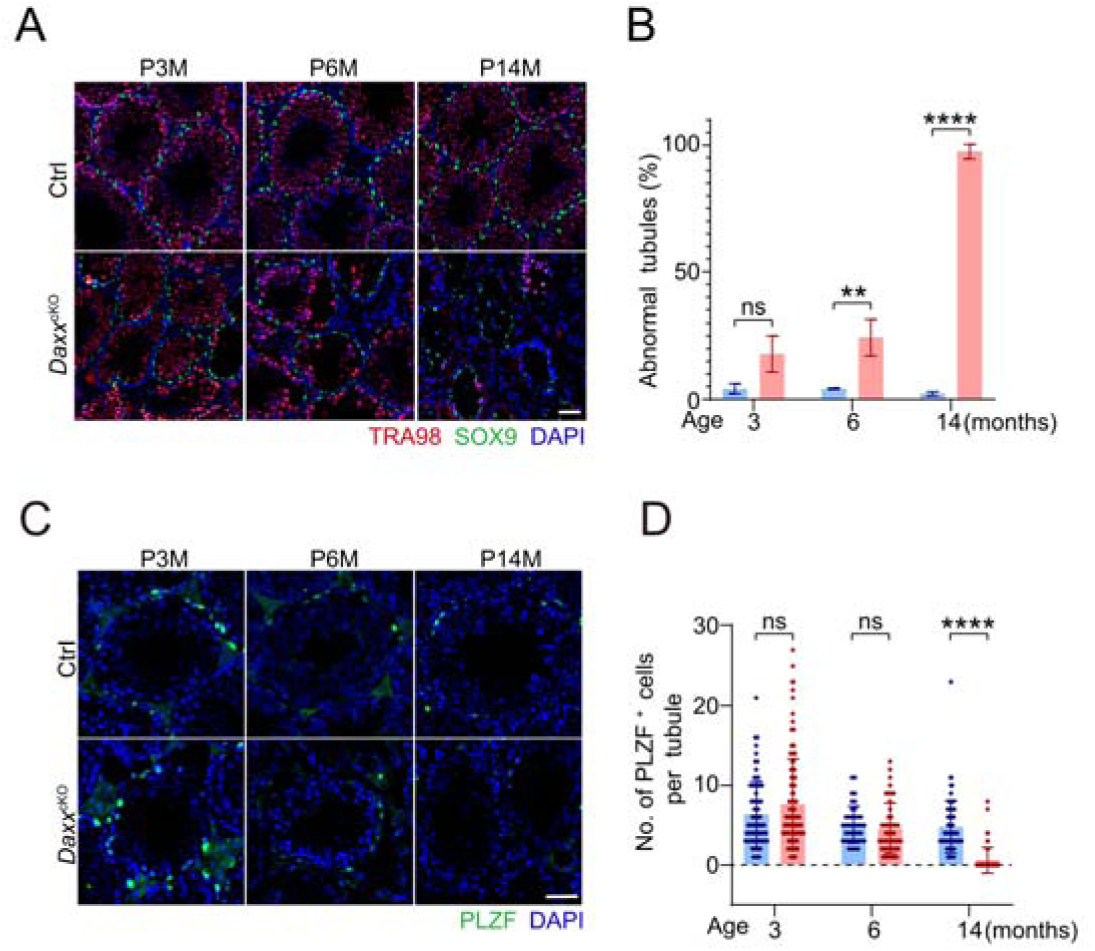
*Daxx* depletion leads to impaired spermatogenesis. Related to Figure 2. (A, B) Immunofluorescence staining of TRA98 (red) and SOX9 (green) indicates that percentage of abnormal seminiferous tubules at 3, 6 and 14 months of age is increased. Scale bar, 50 μm. ns, not significant, \*\**p* < 0.01, \*\*\*\**p* < 0.0001 using two-tailed Student’s unpaired *t*-test. (C, D) Immunofluorescence staining indicates that the number of PLZF-positive cells (green) per tubular cross-section between control and *Daxx*^cKO^ mice at 3, 6 and 14 months of age. Scale bar, 50 μm. ns, not significant, \*\*\*\**p* < 0.0001 using Multiple Mann-Whitney tests.

**Figure S3.**
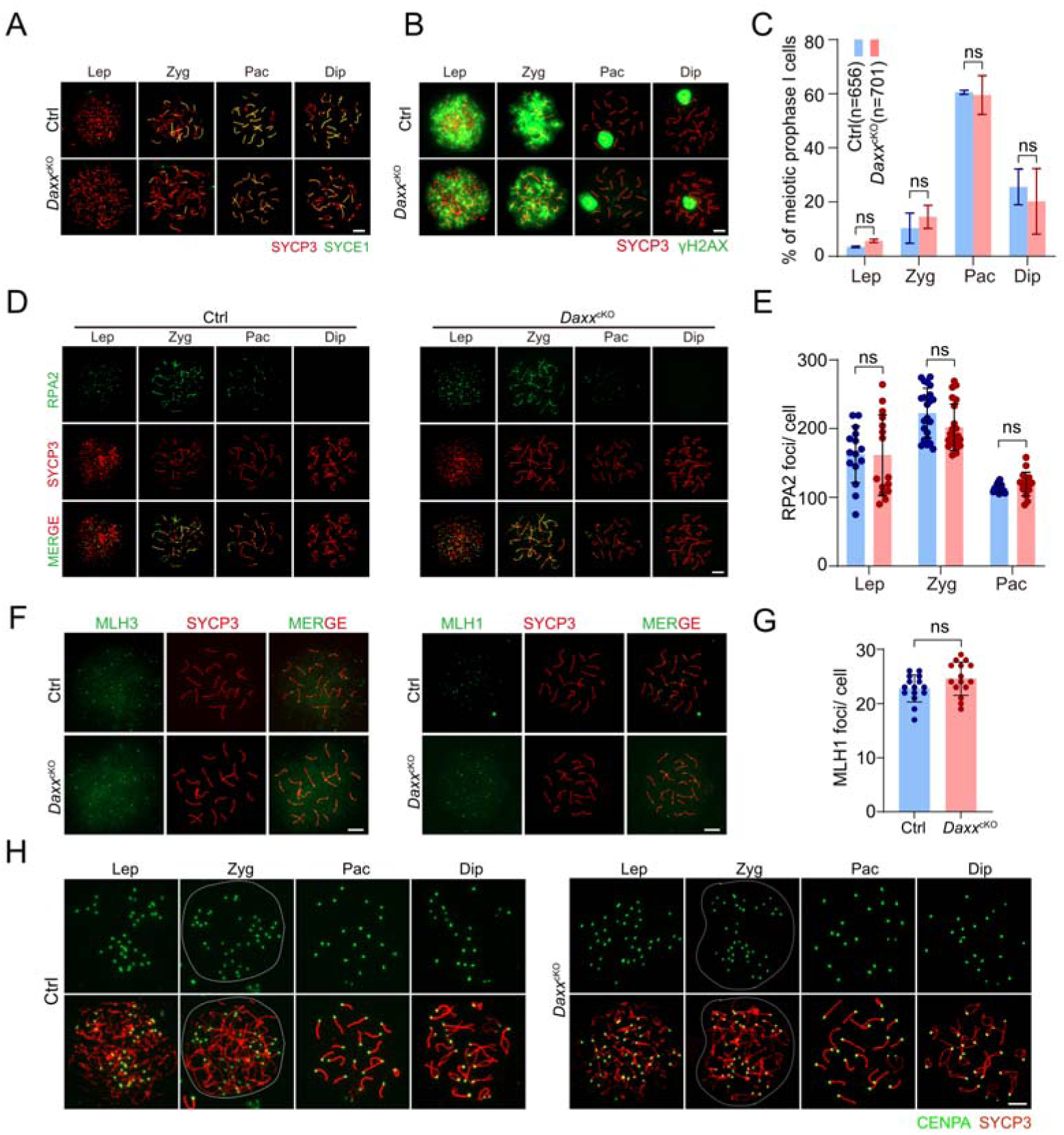
DAXX loss does not affect meiotic chromosomal behaviors. (A) Immunofluorescence of SYCE1 (green) and SYCP3 (red) on chromosome spreads of spermatocytes from control and *Daxx*^cKO^ testes. Scale bar, 10 μm. (B) Immunofluorescence with SYCP3 (red) and γH2AX (green) on chromosome spreads of spermatocytes from control and *Daxx*^cKO^ testes. Scale bar, 10 μm. (C) Percentages of spermatocytes at sub-stages of prophase I in control and *Daxx*^cKO^ mice. ns, not significant. “n” indicates the total number of cells examined from two mice per genotype. (D) Distribution pattern of RPA2 foci on chromatin of spermatocytes from leptotene through diplotene stages in control and *Daxx*^cKO^ mice. Stages were determined by SYCP3 staining. Scale bar, 10 μm. (E) Quantification of focus numbers of RPA2 from WT and *Daxx*^cKO^ mice. Control: n = 15 for lep; n = 22 for zyg; n = 15 for pac; *Daxx*^cKO^: n = 15 for lep; n = 22 for zyg; n = 16 for pac. ns, not significant using Multiple Mann-Whitney tests. (F) Distribution pattern of MLH1 and MLH3 foci on chromatin of spermatocytes at mid-pachytene stages of meiotic prophase I in control and *Daxx*^cKO^ mice. Scale bar, 10 μm. (G) Quantification of focus numbers of MLH1 from WT and *Daxx*^cKO^ mice. Control: n = 15 for steps; *Daxx*^cKO^: n =15. ns, not significant using Multiple Mann-Whitney tests. (H) Immunofluorescence of CENPA (green) and SYCP3 (red) on chromosome spreads of spermatocytes from control and *Daxx*^cKO^ testes. Scale bars, 10 μm. Abbreviation in A-E and H: Lep: leptotene, Zyg: zygotene, Pac: pachytene, and Dip: diplotene.

**Figure S4.**
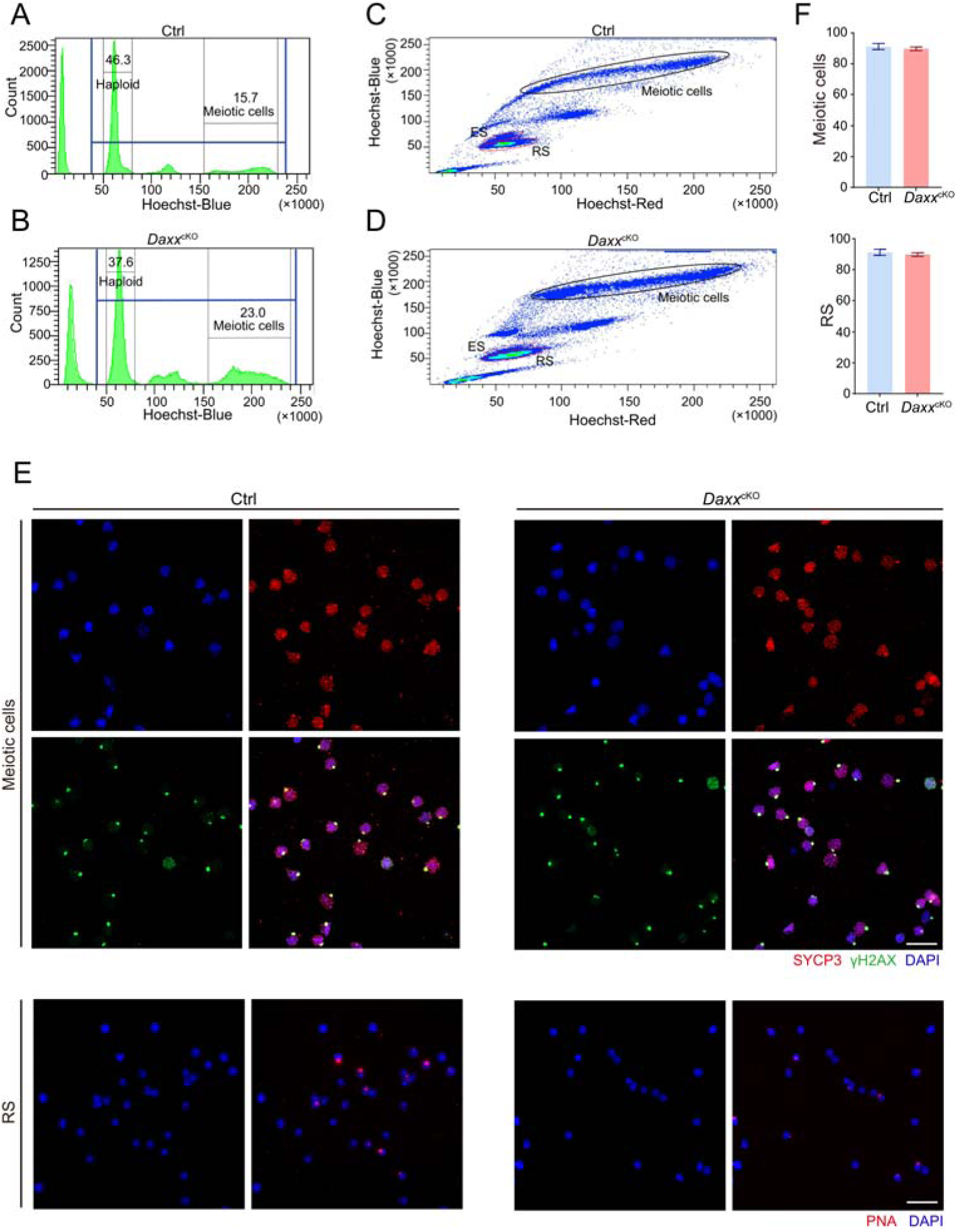
Flow cytometric analysis of testicular cells from control and *Daxx*^cKO^ mice based on Hoechst 33342 fluorochrome and light scattering parameters. Related to Figure 3. (A, B) DNA content exclusion based on Hoechst-blue fluorescence. Cell populations that fall within the blue “DNA” gate include cells with haploid cells and meiosis cells. Low Hoechst-blue events corresponding to debris or dead cells are eliminated. control (A); *Daxx*^cKO^ (B). (C, D) Gating on individual testicular populations based on Hoechst-blue/Hoechst-red fluorescence; RS, round spermatids; ES, elongating spermatids. control (C); *Daxx*^cKO^ (D). (E) Detailed immunofluorescence characterization of sorted meiotic prophase I spermatocytes and RS. Spread nuclei were labeled by staining with γH2AX (green), SYCP3 (red) and DAPI (blue) for meiotic spermatocytes, PNA (red) and DAPI (blue) for RS. Scale bars, 50 μm. (F) Purity statistics of cells obtained by FACS.

**Figure S5.**
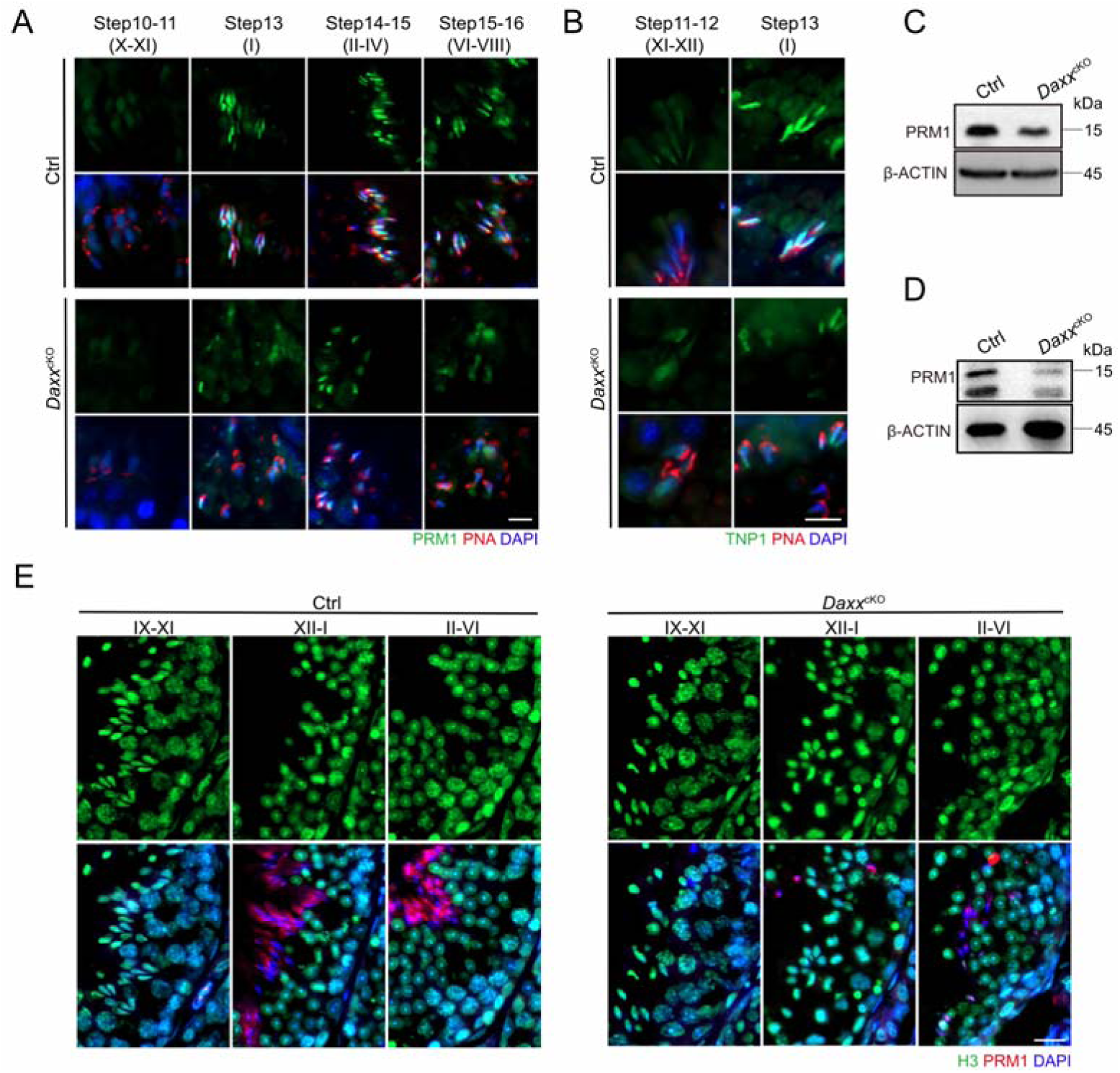
Histone-to-protamine exchange is normal in *Daxx*^cKO^ mice. Related to Figure 3. (A) Representative images of immunofluorescence staining for PRM1 (green), PNA (red), and DAPI (blue) on testicular sections from control and *Daxx*^cKO^ mice. Scale bar, 10 μm. (B) Representative images of immunofluorescence staining for TNP1 (green), PNA (red), and DAPI (blue) on testicular sections from control and *Daxx*^cKO^ mice. Scale bar, 10 μm. (C) Representative western blot showing the protamine protein levels in the testes of control and *Daxx*^cKO^ mice, β-ACTIN serves as the loading control. (D) Representative western blot showing the protamine protein levels in mature sperms of control and *Daxx*^cKO^ mice, β-ACTIN serves as the loading control. (E) Representative images of immunofluorescence staining for H3 (green), PRM1 (red), and DAPI (blue) on testicular sections from control and *Daxx*^cKO^ mice. Scale bar, 20 μm.

**Figure S6.**
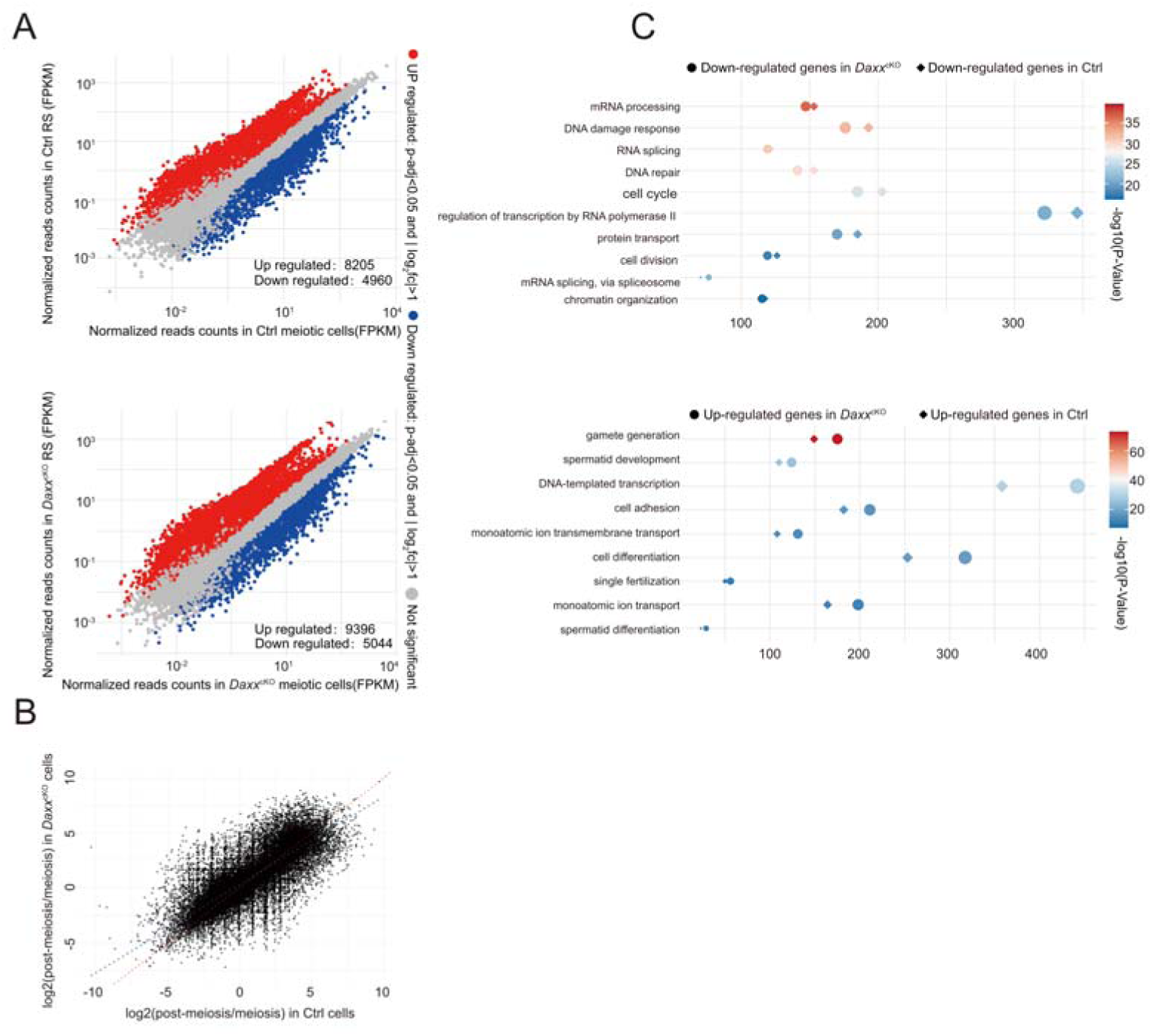
DAXX is implicated in transcriptional repression from meiotic to post-meiotic transition. Related to Figure 5. (A) Scatter plots comparing gene expression profiles of meiotic and post-meiotic cells in the presence or absence of DAXX. (B) Scatter lot of all pairwise log2-fold change in gene expression during meiotic to post-meiotic transition between control and *Daxx*^cKO^ mice. The red dashed line is the Y = X axis and the blue dashed line is the regression line. (C) Functional annotation clustering of down-regulated and up-regulated genes from meiotic to post-meiotic transition in the presence or absence of DAXX.

**Figure S7.**
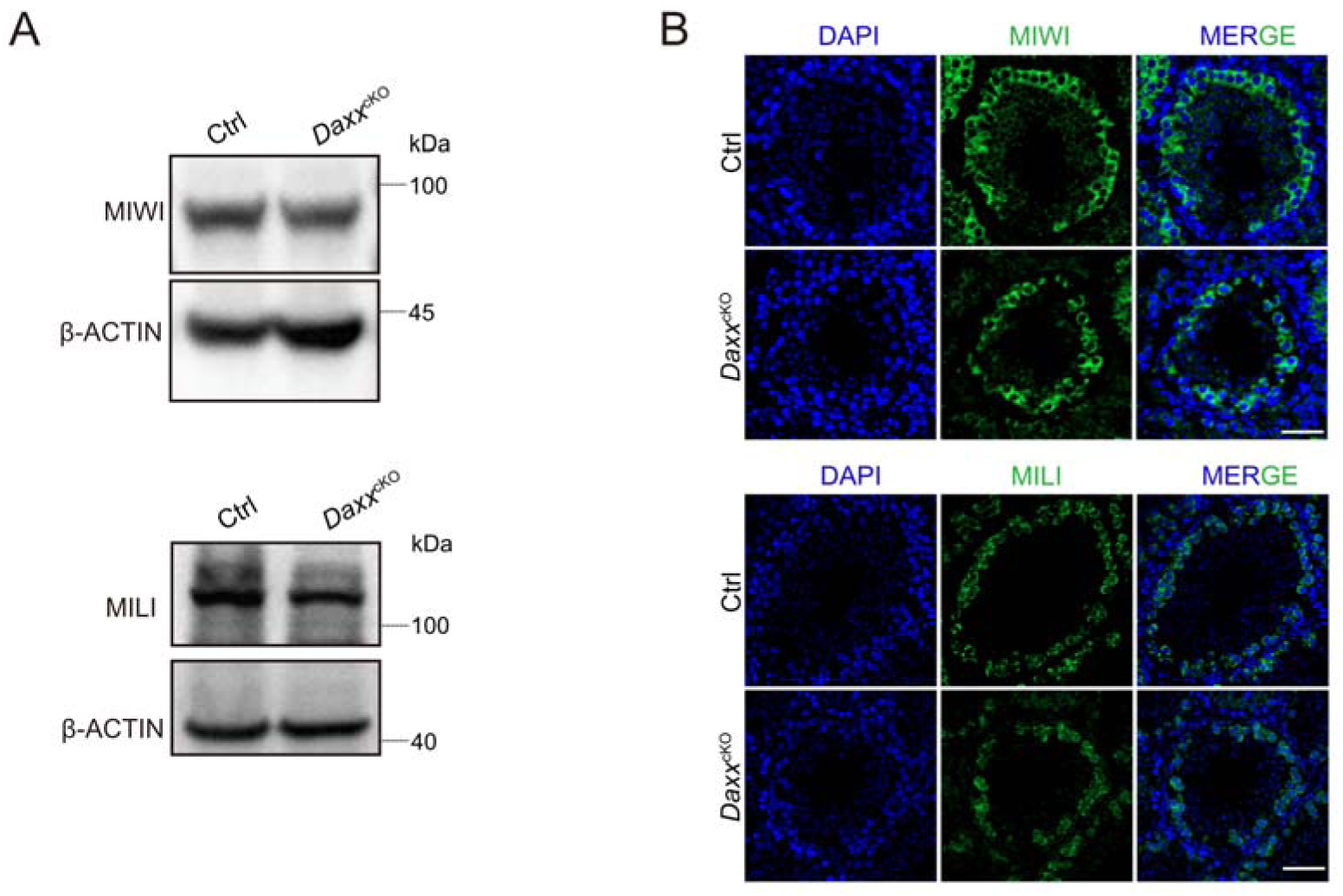
Normal expression of pachytene piRNA-related proteins MIWI and MILI in *Daxx*^cKO^ mice. Related to Figure 6. (A) Western blot of MIWI and MILI in the whole testes of control and *Daxx*^cKO^ mice. β-ACTIN serves as the loading control. (B) Representative images of immunofluorescence staining for MIWI (green) and MILI (green), and DAPI (blue) on testis sections from control and *Daxx*^cKO^ mice. Scale bar, 50 μm.

**Figure S8.**
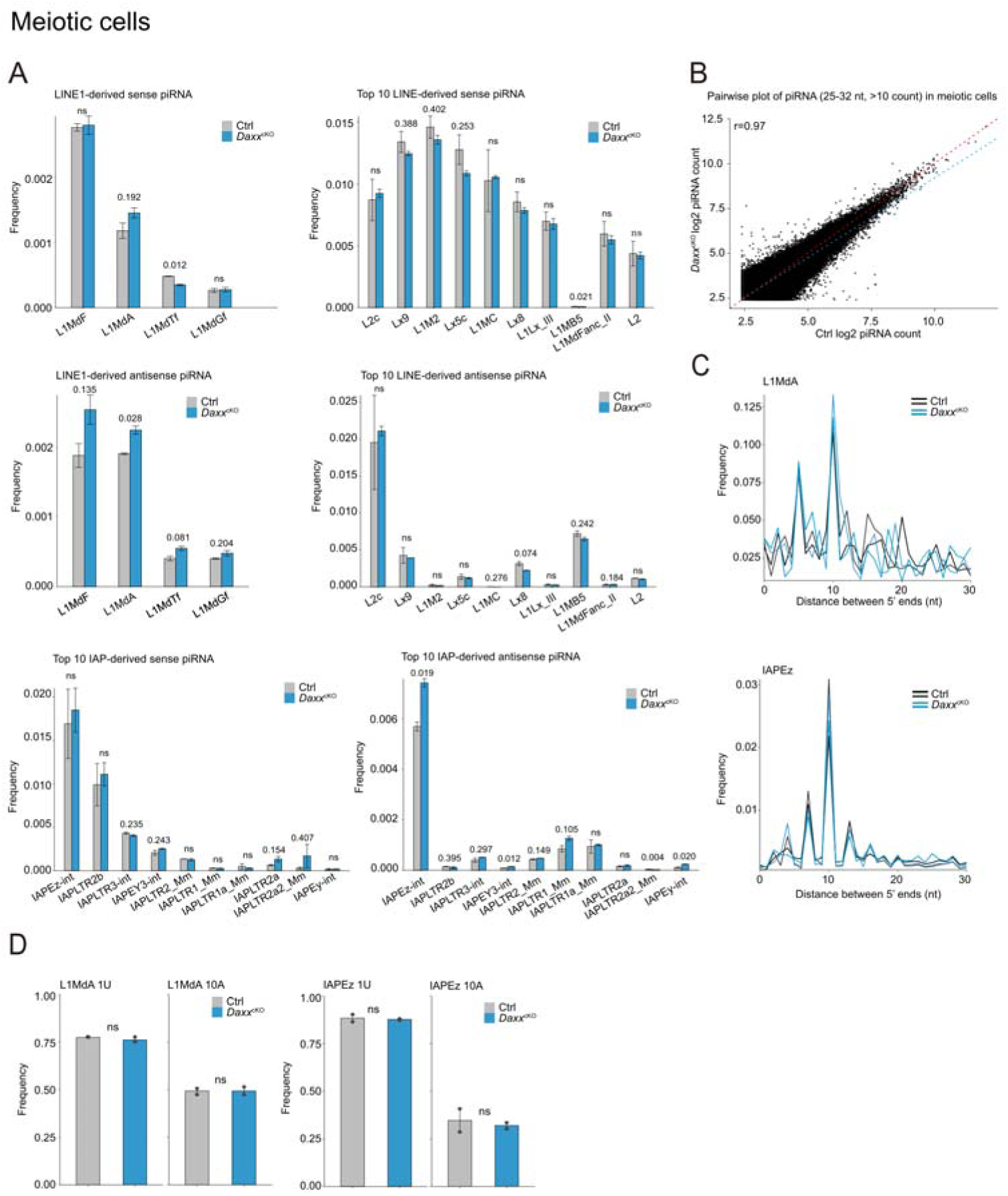
Increased production of piRNAs from L1MdA and normal ping-pong amplification of piRNAs in *Daxx*^cKO^ meiotic cells. Related to Figure 6. (A) Relative frequency of piRNAs mapping to LINE1 and IAP families from control and *Daxx*^cKO^ meiotic spermatocytes. (B) Scatter plots showing mean expression of all piRNAs. The red dashed line is the Y = X axis and the blue dashed line is the regression line. r, Pearson’s correlation coefficient. (C) Ping-pong analysis of piRNAs from control and *Daxx*^cKO^ meiotic spermatocytes: Relative frequency of the distance between 5’ ends of complementary piRNAs mapping to the L1Md_A and IAPEz is shown. (D) Nucleotide features of piRNAs from control and *Daxx*^cKO^ meiotic spermatocytes. Position-specific nucleotide bias analysis of piRNAs demonstrated conserved 1U (position 1 uridine) and 10A (position 10 adenosine) enrichment patterns across annotated genomic elements. Data in A and D, Statistics: Bonferroni-adjusted two-tailed Student’s *t*-tests, with adjusted *p*-values indicated, *p* > 0.5 values are denoted ns (not significant). Data are mean and s.e.m. The original data are available in Table S4.

**Figure S9.**
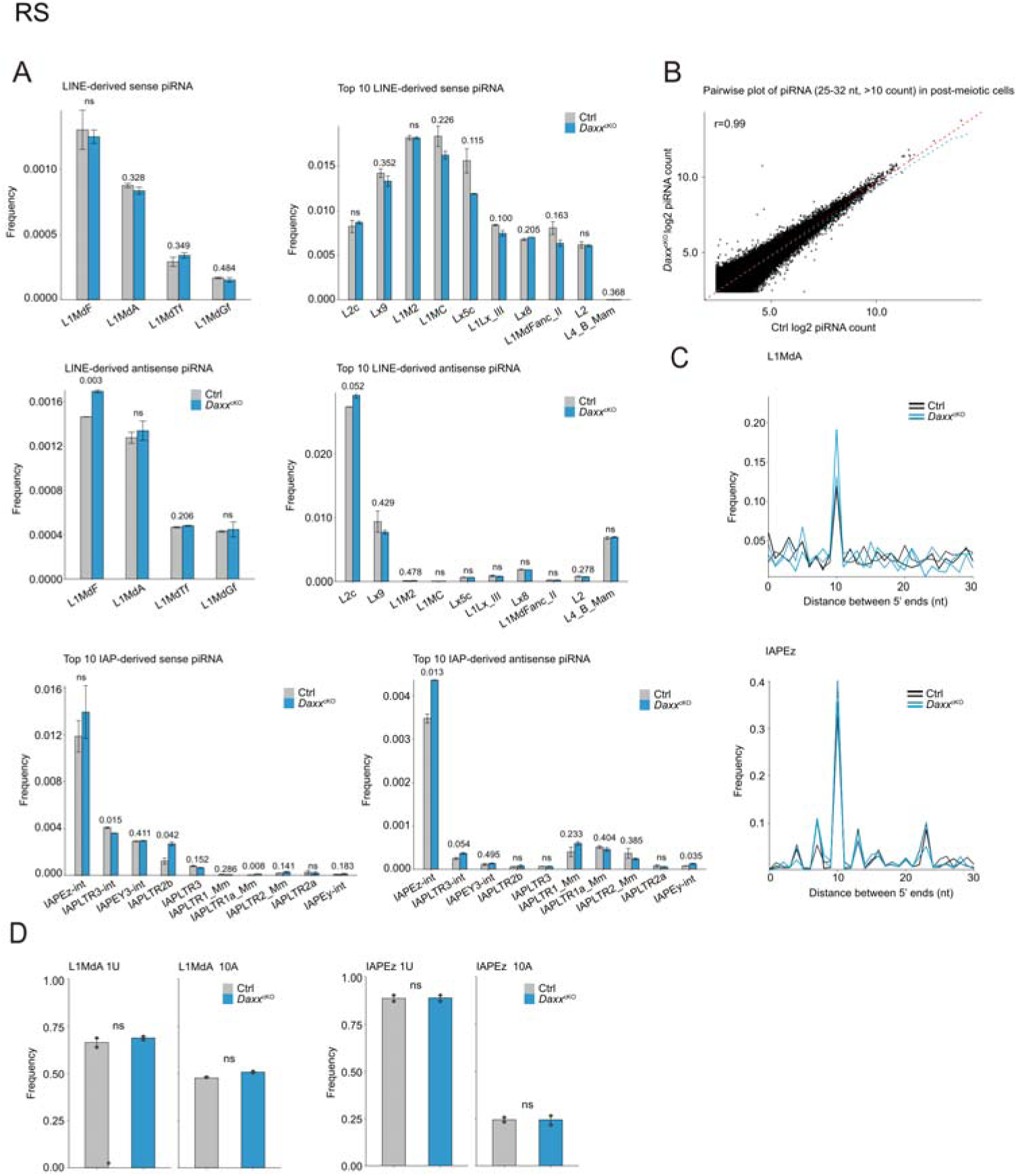
Normal ping-pong amplification of piRNAs in *Daxx*^cKO^ RS. Related to Figure 6. (A) Relative frequency of piRNAs mapping to LINE1 and IAP families from control and *Daxx*^cKO^ RS. (B) Scatter plots showing mean expression of all piRNAs. The red dashed line is the Y = X axis and the blue dashed line is the regression line. r, Pearson’s correlation coefficient. (C) Ping-pong analysis of piRNAs from control and *Daxx*^cKO^ RS. Relative frequency of the distance between 5’ ends of complementary piRNAs mapping to the L1Md_A and IAPEz is shown. (D) Nucleotide features of piRNAs from control and *Daxx*^cKO^ RS. Position-specific nucleotide bias analysis of piRNAs demonstrated conserved 1U (position 1 uridine) and 10A (position 10 adenosine) enrichment patterns across annotated genomic elements. Data in A and D, Statistics: Bonferroni-adjusted two-tailed Student’s *t*-tests, with adjusted *p*-values indicated, *p* > 0.5 values are denoted ns, not significant. Data are presented as mean ± s.e.m. The original data are available in Table S4.

**Figure S10.**
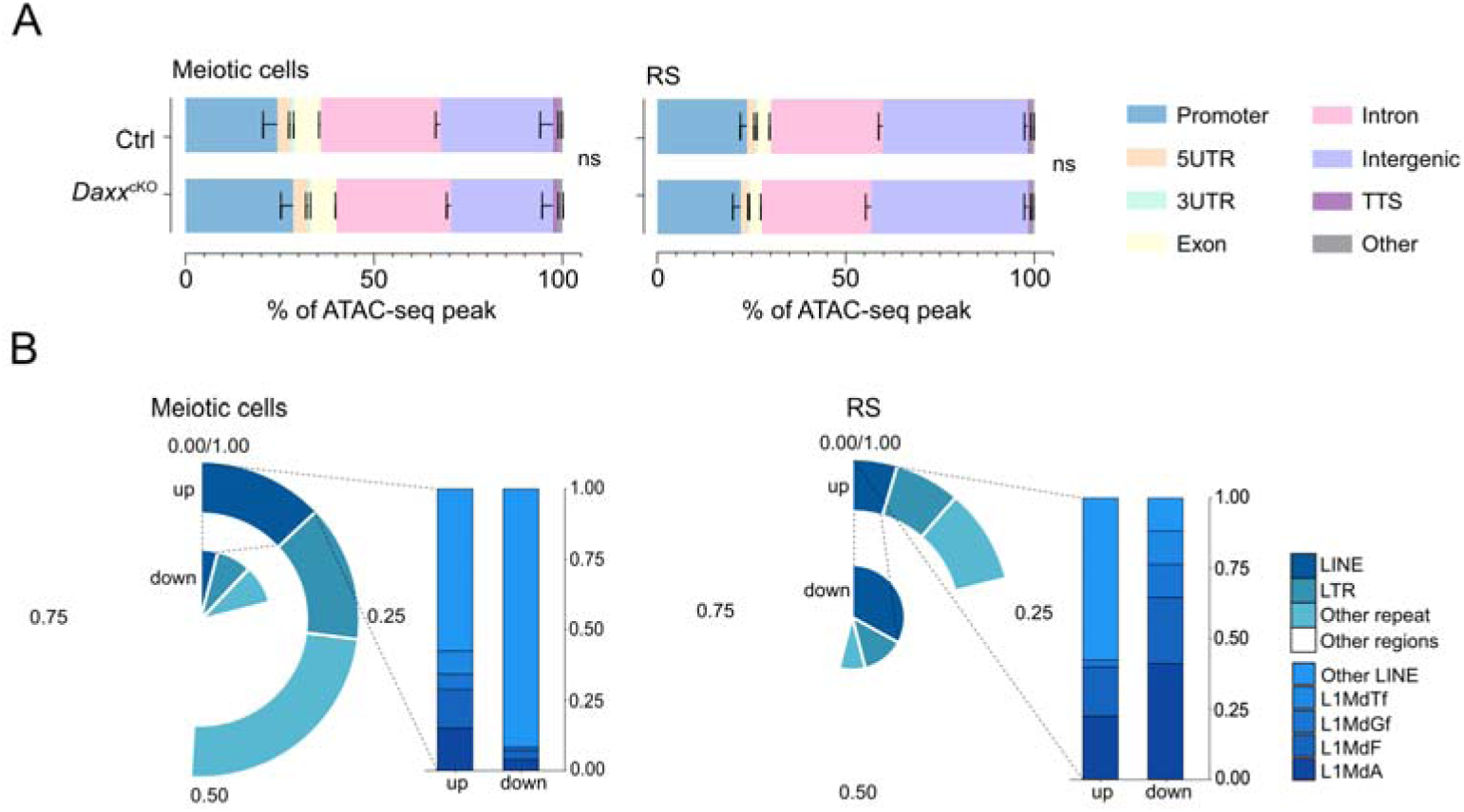
Higher chromatin accessibility at young LINE1 in *Daxx*^cKO^ meiotic cells. (A) Distribution of peaks across different types of functional genomic compartments in control and *Daxx*^cKO^ meiotic cells and RS. (B) Charts show analysis of ATAC-seq peaks with the indicated genomic features. The original data are available in Table S6.

**Figure S11.**
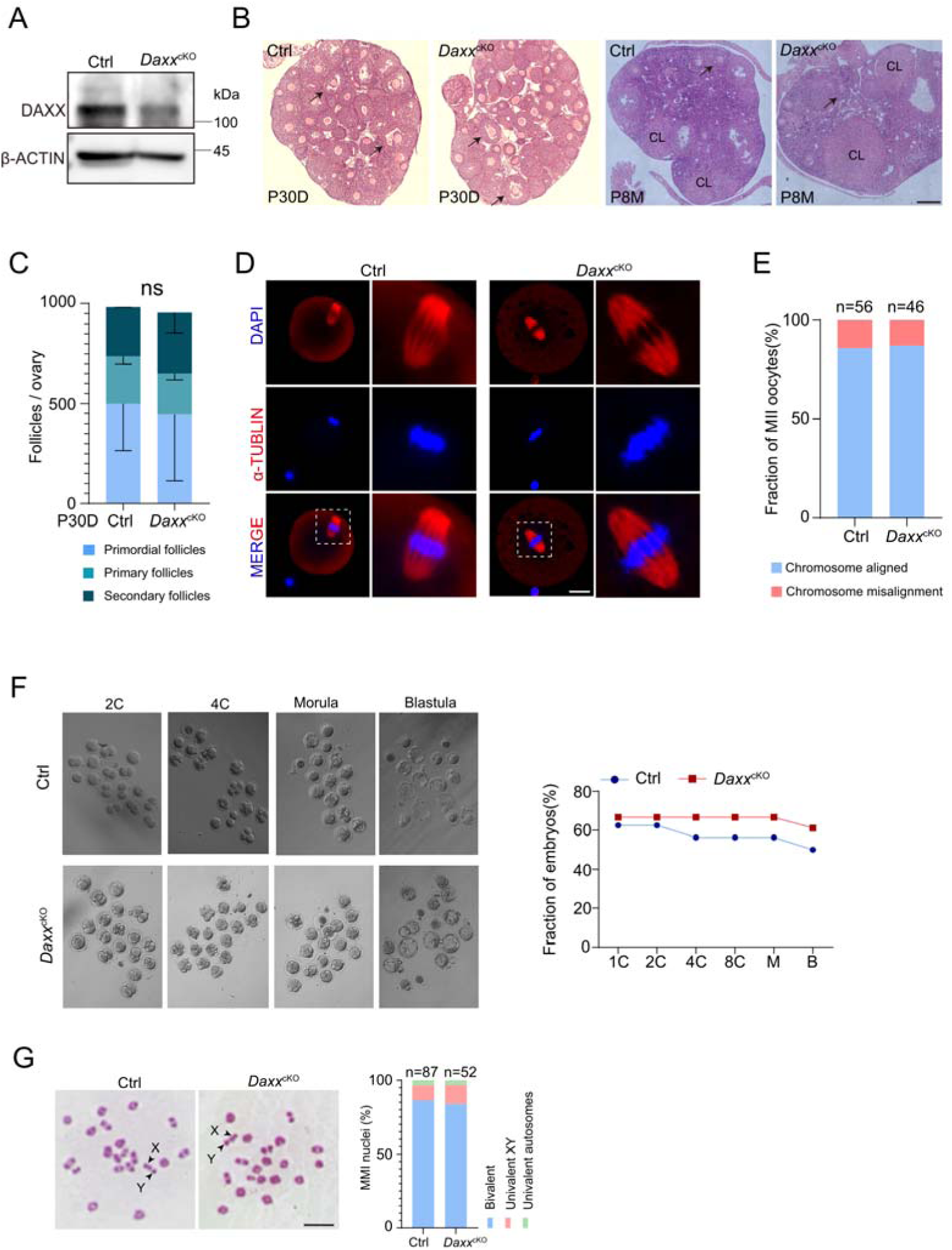
DAXX is dispensable for oogenesis and early embryonic development as well as MMI spermatocytes. (A) Western blot showed that DAXX was deleted in adult *Daxx*^cKO^ ovaries compared to control. β-ACTIN serves as the loading control. (B) Histological analysis of control and *Daxx*^cKO^ ovaries from mice at 1 and 8 months of age. Arrows indicate follicles. CL, corpora lutea. Scale bar, 200 μm. (C) Follicle counts from frozen sections of ovaries at P30D, respectively. Ovaries were stained with YBX2 and follicles counted in every fifth section counts. (D) Analysis of chromosome alignment in MII oocytes. MII oocytes from control and *Daxx*^cKO^ mice were subjected to immunofluorescence labeling with α-tubulin (red) and DNA (blue). Scale bars, 20 μm. (E) Quantification of chromosome alignment in control (n = 56) and *Daxx*^cKO^ (n = 46) MII oocytes. (F) *Daxx*^cKO^ female preimplantation embryos develop normally. IVF was performed with superovulated oocytes from 2-month-old females. Control, 64 oocytes (3 females); *Daxx*^cKO^, 54 oocytes (3 females). Morphology of cultured preimplantation embryos.1C: one-cell embryos, 2C: two-cell embryos, 4C: four-cell embryos, 8C: eight-cell embryos, M: morula, B: blastocyst. (G) Meiotic metaphase I spermatocytes stained with Giemsa and frequencies of nuclei with XY chromosome or autosome dissociation in control (n = 87) and *Daxx*^cKO^ (n = 52) spermatocytes. Chromosomes X and Y are indicated. Scale bars, 10 μm.

## Supplemental table legends

Table S1. List of primers used in this study

<u>Table S1</u>

Table S2. List of differentially expressed genes

<u>Table S2</u>

Table S3. List of differentially expressed transposons

<u>Table S3</u>

Table S4. piRNA analysis

<u>Table S4</u>

Table S5. Whole-genome methylation analysis

<u>Table S5</u>

Table S6. ATAC-seq analysis

<u>Table S6</u>

Table S7. List of reagent or resource

<u>Table S7</u>

